# Ontogenetic modifications produce similar phenotypes in distantly related click beetles (Coleoptera: Elateridae)

**DOI:** 10.1101/2023.03.15.532721

**Authors:** Dominik Kusy, Michal Motyka, Ladislav Bocak

## Abstract

The study analyzes the relationships of click beetles (Elateridae) *Paulusiella* Löbl, 2007, and *Analestesa* Leach, 1824 (=*Cebriognathus* Chobaut, 1899), both incapable of jumping, with soft-bodied habitus, and unknown females. Due to divergent morphology, their positions have been an uncertain issue. We use mitochondrial genomes to test their current placement in Cebrionini (=Cebriognathini) and Elaterinae *incertae sedis*, respectively. We recover *Paulusiella* as a sister to *Hemiops* Laporte, 1838 (Hemiopinae) and *Analestesa* as one of the serially splitting branches in Cardiophorinae, both with robust support. Paulusiellinae **subfam. nov**. is proposed for *Paulusiella. Analestesa* is transferred to Cardiophorinae, and Cebriognathini Paulus, 1981, an earlier synonym of Elaterinae: Cebrionini, is a synonym of Cardiophorinae Candèze, 1859. The click beetles affected by ontogenetic modifications converge to similar forms lacking derived states. As a result, their phylogenetic position cannot be reliably inferred by morphological analyses and needs to be validated by molecular data. Paulusiellinae and *Analestesa* represent two additional cases of the shift to incomplete sclerotization in elaterids raising the total number to six. The present transfers of extant taxa between subfamilies call for a cautious interpretation of morphology in other soft-bodied groups, including the taxa described from amber deposits.

## Introduction

Click beetles (Elateroidea: Elateridae) are well known for their jumping adults (Ribak & Weihs 2011). Still, some click beetles lost their thoracic click mechanism and differ from their relatives in general appearance. Omalisids, drilids (false firefly beetles), and plastocerids were separate families considered the relatives of fireflies, soldier- and net-winged beetles (Crowson 1972, Branham & Wenzel 2003, Lawrence et al. 2011). Only recently, they were included in Elateridae (Bocak et al. 2018, Kusy et al. 2019). Cebrionids (Elaterinae) traditionally contained other non-clicking, soft-bodied elaterids. Although cebrionids were earlier treated as a family (Crowson 1955), they are so slightly modified (Arnett 1949, Rattu 2016) that they were included in Elateridae since applying Hennig’s phylogenetic systematics (Crowson 1981, Bouchard et al. 2011). Later, they were downranked to the tribe Cebrionini in Elaterinae (Kundrata et al. 2014). Still, their classification has never benefited from a molecular study, and the relationships of constituent genera have been poorly corroborated. Concerning the known effect of modified metamorphosis on the phenotype of elateroids (Kusy et al. 2019, 2021), reexamining the phylogenetic position of non-clicking elaterids is needed not only to update the classification but also to understand the evolution of ontogenetic modifications.

Metamorphosis is a crucial innovation that is supposedly connected with the enormous diversity of insects (Nicholson et al. 2014). The transition between larva, pupa and adult is a complex, fine-tuned cascade of steps (Jindra et al. 2015, Jindra 2019) that can be prematurely terminated. Then, some larval and pupal characteristics can persist in adults (Gould 1977, Bocak & Brlik 2008, McMahon & Hayward 2016, Bocek et al. 2019). In the most severely affected elateroids, the imaginal characters are not expressed, and larva-like females are sexually mature (Cicero 2008, Wong 1996, Masek et al. 2014, 2015, Makarov & Kazantsev 2022). The ontogenetic modifications are usually similar within a single group. Still, different levels are known in various lineages (Crowson 1972, McMahon & Hayward 2016).

The phenotypically divergent adults are weakly sclerotized, and never use the clicking mechanism known in their relatives (e.g., drilids, omalisids, cebrionids, etc.). Further, we can encounter shortened or vestigial elytra, sometimes connected with a loss of wings. The abdomen can be larviform (drilids), or at least the ventrites are loose (e.g., Dendrometrinae: Plastocerini; Kusy et al. 2018). The incomplete metamorphosis results in the loss of phylogenetically younger traits in agreement with Baer’s recapitulation law (Løvtrup 1978). The modified soft-bodied forms have been confusing to systematists till an independent source of phylogenetic information became available with the sequencing of DNA. Due to absent apomorphic traits, some affected lineages have been assigned inappropriate high ranks. Alternatively, the unrelated groups were merged into a single taxon (Lawrence et al. 2011, Kazantsev 2013). In such a way, the morphologists defined the earlier superfamily Cantharoidea, and the cantharoid clade in Elateroidea (Crowson 1972, Lawrence 1988, Lawrence et al. 2011).

There are several obstacles that have delayed the studies on modified elaterids. Most neotenic groups are rare compared to their close relatives, and obtaining individuals properly fixed for molecular analyses has not been easy. Additionally, some females often remain unknown. We only estimate that the females are affected by the paedomorphic syndrome from the morphology of males, the relationships, and the absence of females in contrast with numerous males deposited in collections. Wingless females remain in the soil, and some only expose the abdomen during the copulation (observed in *Cebrio*, https://www.youtube.com/watch?v=MlEz6jHCgLo&ab_channel=RobertoLascaroaccessed on Nov. 3, 2022); Bocak et al. 2013, Bocek et al. 2018).

Our study reinvestigates the relationships of the soft-bodied forms placed in the tribe Cebrionini (Elaterinae). We revisit morphological evidence for the earlier proposed concept of cebrionids and look for the morphological traits that could potentially support the molecular relationships. We intend to show that morphologically divergent lineages may converge to similar phenotypes even if they are distantly related. The results affect the Linnean classification of investigated taxa. We believe that revising the traditional placement of extant soft-bodied elaterids might contribute to a better understanding of morphological evolution. We expect, analogically to click beetle phylogeny, further changes in the placement of other strongly modified groups. The difficulties with classifying extant soft-bodied forms should also be considered in works on fossils for which only morphological and often incomplete data are available.

## Methods

### The compilation of the dataset

The mitogenomic dataset was assembled from newly sequenced mitogenomes of *Cebrio* sp., *Cebriorhipis* sp., *Analestesa arabica* (Paulus, 1981), *Paulusiella serraticornis* (Paulus, 1972), *Quasimus* sp., and *Hemiops* sp. The voucher numbers and complete locality data are reported in Tab.

1. Further mitogenomic data were taken from the dataset published by Kusy et al. (2021).

### Laboratory procedures, data handling, and morphological investigation

The total DNA was isolated from alcohol-preserved or dry-mounted samples. The vouchers were used for morphological investigation. They were dissected after short relaxation in 50% ethanol. The structures were treated in hot 10% KOH for a short time. The photographs were taken by a Canon M6 Mark II camera attached to an Olympus SZX16 binocular microscope. Stacks were assembled using Helicon Focus software and processed in Photoshop 6.0. Vouchers are deposited in the collections of the collectors and of Biodiversity & Molecular Evolution at CATRIN, Olomouc.

DNA was extracted using Qiagen MagAttract HMW DNA extraction kit, and eluted in AE buffer. Short insert size library constructions (∼320 bp) and subsequent paired-end (2 × 150 bp) sequencing of the samples were done by Novogene, Inc., Beijing, using Illumina NovaSeq 6000. Raw Illumina reads were quality checked with FastQC and filtered with fastp 0.21.0 (Chen et al. 2018) using -q 28 -u 50 -n 15 -l 50 settings. Filtered reads were used for final mitogenome assemblies. The mitogenomes were built de novo using the NOVOPlasty v.2.7.2 pipeline (Dierckxsens et al. 2017). NOVOPlasty was run with the default settings except the kmer value when we used a multi-kmer strategy with the following kmer sizes of 25, 39, 45, and 51. We used as seed the single fragment of *Oxynopterus* sp. cox1 gene available in GenBank (HQ333982). In the case of *Quasimus* sp., the mitochondrial fragments were mined from unpublished transcriptomic data that were mapped on the mitochondrial genome of *Cardiophorus signatus*, and manually curated in Geneious v.7.1.9. The newly assembled mitochondrial genomes were annotated using the MITOS2 webserver (Bernt et al. 2013) with the invertebrate genetic code and RefSeq 63 metazoa reference. The annotation, circularization, and start + stop codons corrections of protein-coding genes (PCSGs) were performed manually in Geneious v.7.1.9. The sequences of newly produced mitochondrial genomes were deposited into the Mendeley database DOI:10.17632/73dmw4czm3.1.

### Phylogenetic analyses

The six mitochondrial genomes of click beetles were merged for the purpose of phylogenetic analyses with thirty earlier published mitochondrial genomes (36 ingroup taxa and 1 outgroup, Kusy et al. 2021). The dataset contained terminals belonging to ten subfamilies of Elateridae. The nucleotide sequences of protein-coding genes (PCG) were aligned using TransAlign (Bininda-Emonds 2005). In addition, nucleotide sequences of rRNA genes and translated amino acid sequences of PCGs were aligned with Mafft v.7.407 using the L-INS-i algorithm (Katoh & Standley, 2013). The aligned data were concatenated with FASconCAT-G v.1.04 (Kück & Longo 2014). We compiled the following datasets: (A) 13 PCG mtDNA and 2 rRNA mtDNA genes partitioned by gene or unpartitioned; (B) 13 mitochondrial PCGs and by gene or unpartitioned; (C) 13 mitochondrial PCGs masked by degen software (Steenwyk et al. 2020) partitioned by a gene or unpartitioned; (D) amino acid level analysis of 13 mitochondrial PCGs. The degree of missing data and overall pairwise completeness scores across all datasets was inspected using AliStat v.1.7. (Thomas et al. 2020) (Supp. Fig. S1).

Phylogenetic inferences were performed under maximum likelihood (ML) optimization using IQ-Tree2.1.2 (Minh et al. 2020), and Bayesian inference (BI) using PhyloBayes MPI v.1.8 (Lartillot et al. 2013). Before ML tree searches, best-fitting model selection for each partition was performed with ModelFinder (Chernomor et al. 2016, Kalyaanamoorthy et al. 2017) using the -MFP. All datasets were tested against a complete list of models. We used the edge-linked partitioned model for tree reconstructions (-spp option) allowing each partition to have its own rate.

Ultrafast bootstrap values (Hoang et al. 2018) were calculated for each tree using -bb 5000 option. In the PhyloBayes analysis, unpartitioned datasets A, and D were analyzed under the site-heterogeneous mixture CAT + GTR + Γ4 model for all searches. Two independent Markov chain Monte Carlo (MCMC) were run for each dataset. We checked for the convergence in the tree space with bpcomp program and generated output of the largest (maxdiff) and mean (meandiff) discrepancy observed across all bipartitions and generated a majority-rule consensus tree using a burn-in of 30% and sub-sampling every 10th tree. Additionally, we used the program tracecomp to check for convergence of the continuous parameters of the model.

We employed several tests to investigate alternative phylogenetic relationships, including the approximately unbiased AU-test (Shimodaira 2002), the p-SH (p-value of the Shimodaira-Hasegawa test) (Shimodaira & Hasegawa 1999), the KH-test (one-sided Kishino-Hasegawa test) (Kishino & Hasegawa 1989), the p-WKH (p-value of weighted KH test), the p-WSH (p-value of weighted SH test), c-ELW (Expected Likelihood Weight) (Strimmer & Rambaut 2002), and bp-RELL (bootstrap proportion using RELL method) (Kishino et al. 1990). To evaluate the hypothesized placement of focal taxa (*Analestesa arabica, Cebriorhipis* sp., and *Paulusiella serraticornis*) within Elaterinae, we tested the maximum likelihood (ML) tree topology against three alternative topologies suggested by earlier classifications: (A) *Analestesa arabica* as sister to *Cebrio* and *Cebriorhipis*, (B) *Paulusiella serraticornis* as sister to *Cebrio* and *Cebriorhipis*, and (C) the clade containing all focal taxa. IQ-TREE2 (Minh et al. 2020) was used to perform all tests, with per-site log-likelihoods calculated using the --test-weight --test-au --sitelh parameters and 10,000 replications

## Results

### Molecular phylogeny

The Bayesian and maximum likelihood mitogenomic analyses indicate high support for the polyphyly of Cebrionini in the traditional sense (Figs. 1A, B; Supp. Figs. S2-S11). Only *Cebrio* sp. and *Cebriorhipis* sp. are members of Elaterinae (BS 99%, PP 0.97). *Paulusiella serraticornis* was regularly a sister to *Hemiops* sp. (BS 100%, PP 1.00). Still, the clade was variably a sister to the remaining elaterid subfamilies or a sister to the non-Elaterinae subfamilies (Fig. 1A, B, Supp. Figs. S2–S11). *Analestena arabica* was firmly placed within Cardiophorinae (BS 100%, PP 1.00)) as the second split following *Globothorax femoralis*. The position of Negastriinae as a putative sister of Cardiophorinae was recovered only by some analyses (Supp. Figs. S5, S6, S8). Alternatively, the genus was found as a sister to the Agrypninae (Fig. 1A), or the Cardiophorinae + Agrypninae clade (Fig. 1B). No analysis suggested alternative positions for the focal taxa. The tests rejected the relationships of *Paulusiella* and *Analestesa* with Cebrionini (*Cebriorhipis* and *Cebrio*; Elaterinae; Tab. 2). We separately considered both genera in Cebrionini, and alternatively, either of them as a sister to *Cebrio* and *Cebriorhipis* (Tab. 2). We did not test the relationships between subfamilies and the monophyly of Elateridae as the dataset does not provide enough support for the deepest relationships.

**Figure 1.**
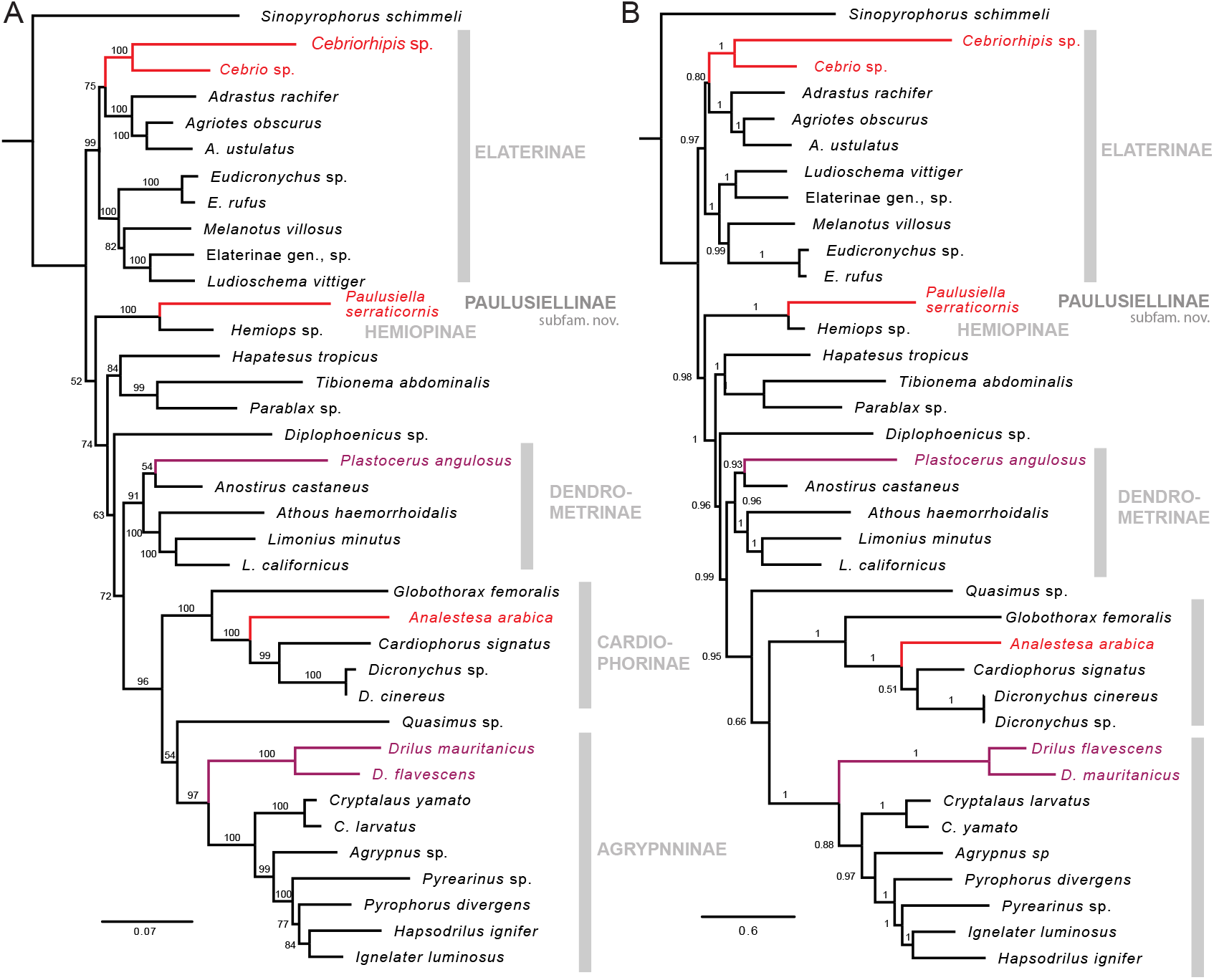
The mitogenomic relationships of soft-bodied and/or non-clicking genera *Paulusiella, Cebrio, Scaptolenus, Analestesa, Plastocerus, Drilus*, and their clicking relatives. The numbers at branches designate bootstrap values and posterior probabilities. A – the maximum likelihood analysis of thirteen protein-coding mitochondrial genes at the nucleotide level with coding masked by the degen software and partitioned by genes. B – The Bayesian analysis using PhyloBayes at the nucleotide level. The trees with full-length branches and the results of additional analyses are shown in Supp. Figs. S2–S11. Red – the taxa earlier placed in Cebrionini; magenta – non-clicking elaterids earlier placed in families Drilidae and Plastoceridae.

**Table 1.** The list of newly sequenced samples. For the list of publicly available samples, see Supp. Tab. S1.

| Taxon | Voucher | Geographic origin |
| --- | --- | --- |
| *Paulusiella serraticornis* | G21004 | Iran, Kerman prov., Gebal Barez mts., 1345 m, 26 km N of Jiroft, wadi, 28°54'14"N 57°40'32"E, 27. May 2018, Vit Kuban leg., coll. V. Kuban. |
| *Analestesa arabica* | G21003 | Kingdom of Saudi Arabia, Hieth, 2. May 1975, 40 km S of Riyadh, 24°18'9", 46°42'27", W. Büttiker leg., coll. L. Bocak. |
| *Cebrio* sp. | G19010 | Italy, Sardinia, 2 km W of Irgutosu, 62 m, 39°31'31"N, 8°28'20"E, D. Ahrens & S. Fabrizzi leg., coll. L. Bocak. |
| *Cebriorhipis* sp. | G22001 | Indonesia, Bali, Tamblingan Lake, 1000–1300 m, 8°15'33"S, 115°6'14"E. 2.–17. Feb 2004, S. Jakl leg., coll. D. Kusy. |
| *Quasimus* sp. | R19010 | Japan, Kunimidake, 35°27'3 "N, 136°21'37"E, 14. May 2015, T. Sota leg., coll. L. Bocak. |
| *Hemiops* sp. | G19002 | Malaysia, Perak, km 24 Rd Tapah-Ringlet, 230 m, 4°18'39"N, 101°19'52"E, 19. Apr. 2013, L. Dembicky leg., coll. L. Dembicky. |

**Table 2.**
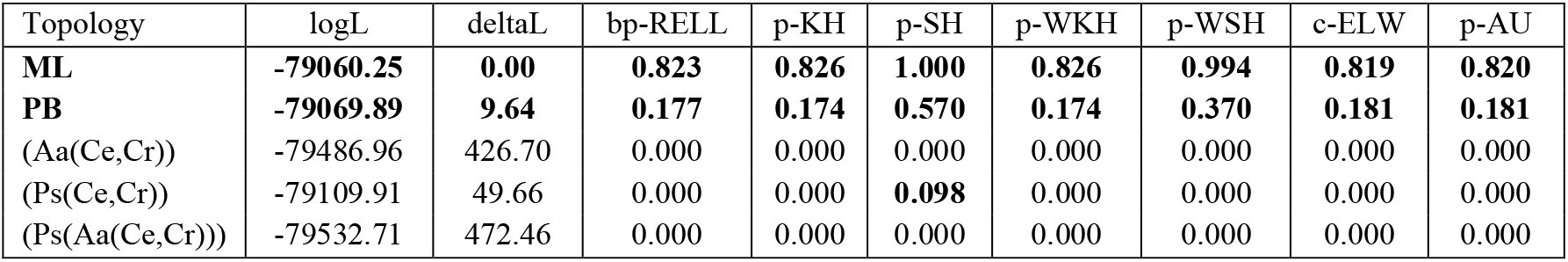
Results of the alternative tree topologies likelihood testing. Aa: *Analestesa arabica*; Ce: *Cebrio* sp.; Cr: *Cebriorhipis* sp.; Ps: *Paulusiella serraticornis;* deltaL: logL difference from the maximal logL in the set; bp-RELL: bootstrap proportion using RELL method; p-KH: p-value of one-sided Kishino–Hasegawa); p-SH: p-value of Shimodaira–Hasegawa test; p-WKH: p-value of weighted KH test; p-WSH: p-value of weighted SH test; c-ELW: Expected Likelihood Weight; p-AU: p-value of approximately unbiased (AU) test. Bold text represents the accepted test.

| Topology | logL | deltaL | bp-RELL | p-KH | p-SH | p-WKH | p-WSH | c-ELW | p-AU |
| --- | --- | --- | --- | --- | --- | --- | --- | --- | --- |
| **ML** | **-79060.25** | **0.00** | **0.823** | **0.826** | **1.000** | **0.826** | **0.994** | **0.819** | **0.820** |
| **PB** | **-79069.89** | **9.64** | **0.177** | **0.174** | **0.570** | **0.174** | **0.370** | **0.181** | **0.181** |
| (Aa(Ce,Cr)) | -79486.96 | 426.70 | 0.000 | 0.000 | 0.000 | 0.000 | 0.000 | 0.000 | 0.000 |
| (Ps(Ce,Cr)) | -79109.91 | 49.66 | 0.000 | 0.000 | **0.098** | 0.000 | 0.000 | 0.000 | 0.000 |
| (Ps(Aa(Ce,Cr))) | -79532.71 | 472.46 | 0.000 | 0.000 | 0.000 | 0.000 | 0.000 | 0.000 | 0.000 |

### Morphology

#### Paulusiella serraticornis (Paulus, 1972)

*Escalerina serraticornis* Paulus, 1972: 38 (in Karumiidae; now Dascillidae: Karumiinae). (Figs. 2A–P)

**Fig. 2.**
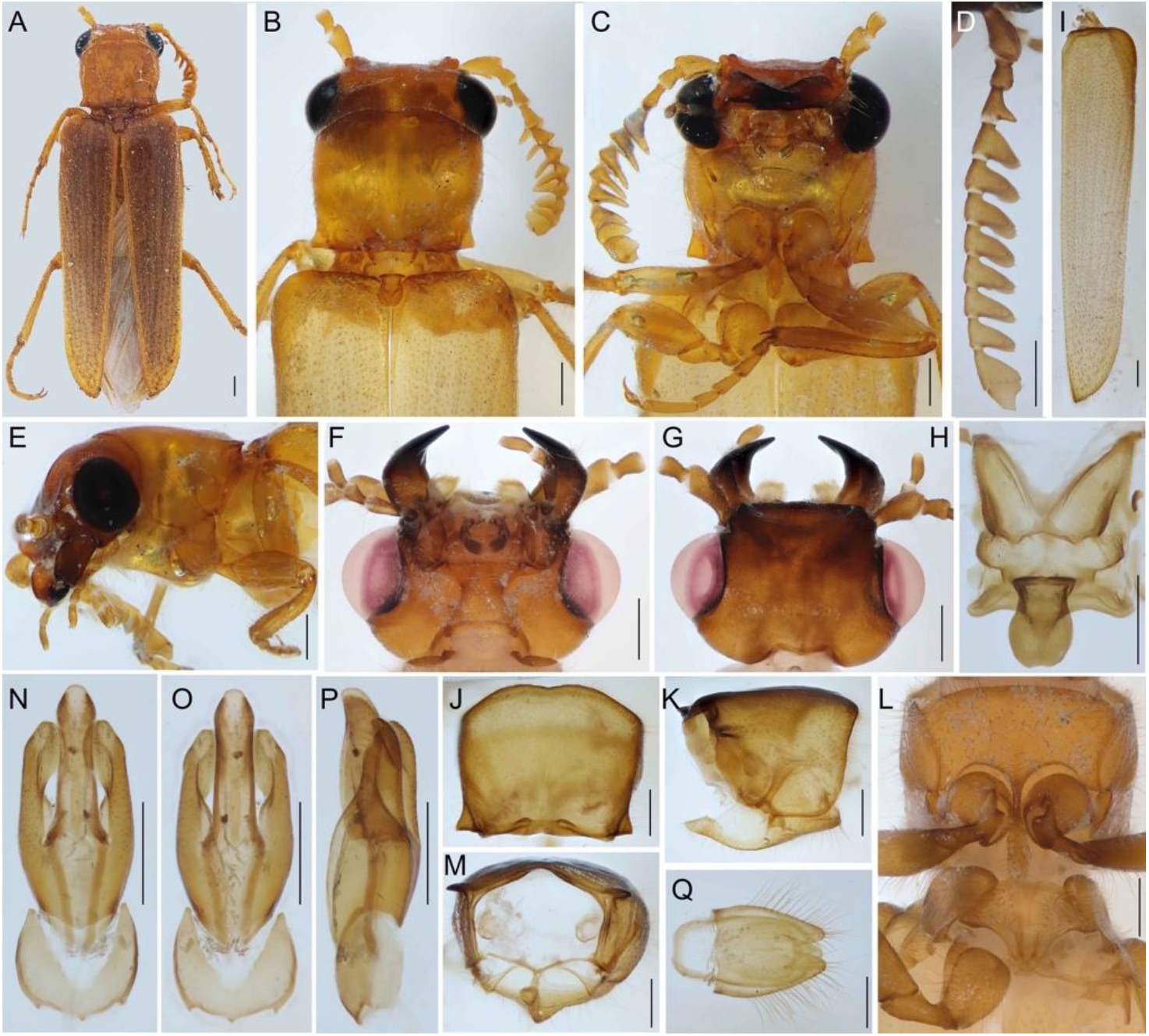
*Paulusiella serraticornis* from Iran. A – general appearance, dorsal view; B, C – frontal part of the body, dorsally and ventrally; D – antenna; E – head and pronotum in lateral view; F, G – head, ventral and dorsal view; H – mesonotum; I – elytron; J–L –pronotum, dorsally, laterally, posterior view, M prosternal process, and mesosternal pit; N–P male genitalia, ventrally, dorsally, laterally; Q terminal abdominal segments. Scales 0.5 mm.

#### Redescription

Male. Body 5–7 mm long, slender, slightly flattened, light brown colored, vestiture of upper surfaces with erect, long bristles (Figs. 2A–C).

Head slightly declined, transverse, with small antennal sockets, including eyes greater than prothoracic width, gradually narrowed posteriorly. Frons and vertex flat, with lateral carinae behind antennal pits, without ocelli (Figs. 2F, G). Eyes protuberant, rounded, finely facetted. Antennal insertions widely separated, covered by protuberant edge from above (Fig. 2B). Anterior edge of clypeus straight (Fig. 2C), gular sutures narrowly separated, and cervical sclerites well-sclerotized (Fig. 2F). Antennae reaching elytral humeri, antennomere 1 robust, long, antennomere 2 small, but longer than width, antennomeres 4-10 flabellate, terminal antennomere flat (Fig. 2D). Labrum concealed beneath clypeus; mandibles robust, curved, with dorso- and ventrolateral edges (Fig. 2E).

#### Mandibular apex

unidentate, incisor edge simple, without mola (Figs. 2F, G), maxilla with setose mala; maxillary palpi slender, 4-segmented, apical palpomere cylindrical; labium tiny, labial palpi 3-segmented, cylindrical (Fig. 2F).

Prothorax transverse (Fig. 2B), pronotum without carinae, maximum width 1.23 times length, widest in anterior third, only slightly narrower at base, sides sinuate (Figs. 2B, J). Prothorax basally narrower than elytral bases; lateral pronotal carina visible posteriorly. Posterior angles of pronotum strongly acute (Fig. 2K). Posterior edge of pronotum sinuate. Prosternum about as long as prosternal process, process long slender, edge curved in lateral view (Figs. 2C, K); apex of prosternal process does not reach mesosternal pit (Fig. 2L); promesothoracic clicking mechanism non-functional; procoxal cavities open, narrowly separated (Fig. 2C). Elytra cover whole abdomen, tapering to apex, widest at humeri, without apparent costae or rows of punctures (Fig. 2I). Elytra free apically (Fig. 2A), Elytral pleuron very short. Scutellum well developed; abruptly elevated; anteriorly straight, posteriorly broadly rounded and surpassing elytral surface, anterior edge of mesoventrite rounded (Fig. 2H), meso- and meta coxal cavities narrowly separated. Metasternum long, discrimen complete, posterior transverse suture apparent, posterior margin deeply emarginate between coxae. Hind wing present.

Legs slightly compressed, with long, gradually widened trochanters, femoral attachments oblique; femora twice wider than tibiae (Fig. 2C), tibiae with simple outer edge and two long apical spines; tarsomeres slender, five segmented, without ventral pads, claws paired, long, slender, and simple.

Abdomen with six visible abdominal ventrites, ventrite 1 divided by metacoxae, ventrite 2 without process; penultimate tergite deeply emarginate, ultimate tergite very small, narrow (Fig. 2Q). Male genitalia trilobate; symmetrical. Phallus stout, short, basally merged with parameres (Figs. 2N– P).

Females of all species unknown.

#### Biology, distribution and species diversity

All species have been reported from semidesert ecosystems of southwestern Asia, and the highest diversity is known from Iran. Males are commonly collected at the light. The biology is unknown, and the larvae and females are presumably endogenous. The mite species *Trochometridium kermanicum* Mortazavi & Hajiqanbar, 2011 was found on *Paulusiella* sp. in Iran (Mortazavi et al. 2011).

The genus contains six species: *P. serraticornis* (Paulus, 1972) (Iran); *P. richteri* (Mandl, 1974) (Iran); *P. fossulatipennis* (Mandl, 1974) (Pakistan); *P. pallida* (Mandl, 1974) (Iran); *P. holzschuhi* (Mandl, 1979) (Iran); *P. sweihana* (Geisthardt, 2009) (United Arab Emirates).

#### Analestesa arabica (Paulus, 1981)

*Cebriognathus arabicus* Paulus, 1981: 261.

(Figs. 3A–P)

**Fig. 3.**
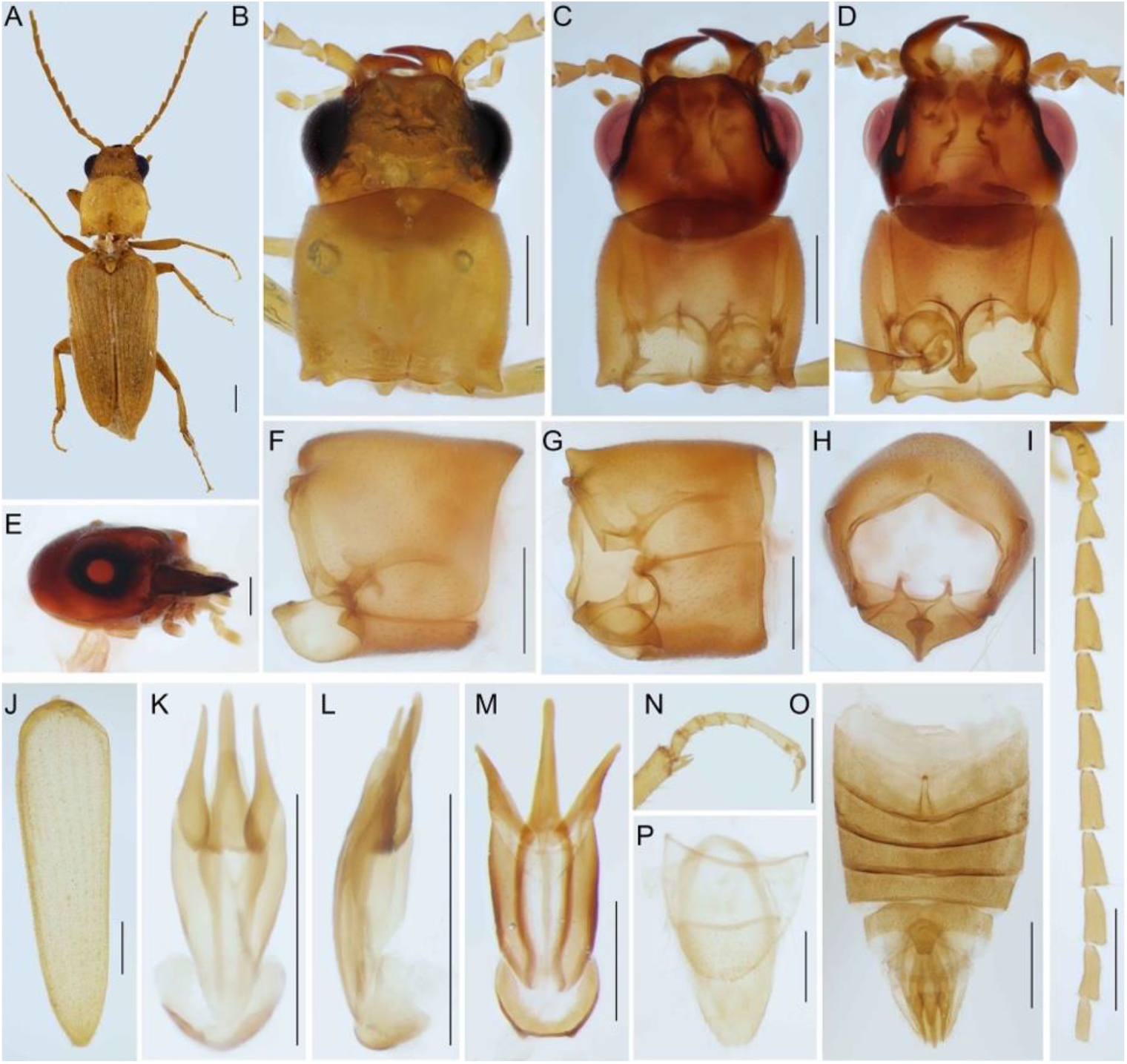
*Analestesa arabica* (Paulus, 1981) from Saudi Arabia (except for Fig. 3I). A – general appearance, dorsal view; B–D – frontal part of the body, dorsally (B, C) and ventrally); E – head in lateral view; F–H – pronotum, lateral, dorsolateral, and posterior view; I – antenna; J – elytron; K, M male genitalia, ventrally, dorsally, laterally; l – Male genitalia of *Dicronychus cinereus* (Herbst, 1784); N – hind tarsus and apical part of tibia; O – abdomen, ventral view; P – terminal abdominal segments. Scales 0.5 mm.

#### Redescription

Male. Body 6 mm long, slender, slightly flattened, light brown colored, vestiture dense and short (Fig. 3A).

Head prognathous, slightly transverse, including eyes equals prothoracic width. gradually narrowed posteriorly; dorsally flat, without ocelli (Figs. 3B–D). Eyes slightly protuberant, rounded, finely facetted. Antennal insertions widely separated, covered by protuberant edge from above (Figs. 3B, C, E); clypeus concave (Fig. 3C), gular sutures narrowly separated, and cervical sclerites sclerotized (Fig. 3D). Antennae reaching mid of elytra, antennomere 1 robust, long, antennomere 2 small, antennomeres 4–11 filiform, terminal antennomere shorter than preceding one (Fig. 3I).

Labrum concealed beneath clypeus; mandibles robust, abruptly curved (Fig. 3E). Mandibular apex and incisor edge simple, maxilla with setose mala; maxillary palpi slender, 4-segmented, apical palpomere parallel-sided; labium tiny, labial palpi 3-segmented (Fig. 3D).

Prothorax transverse, pronotum without carinae, maximum width 1.18 times length, widest in middle, sides convex (Fig. 3B). Prothorax basally narrower than elytral bases; lateral pronotal edge absent (Figs. 3F–H). Posterior angles of pronotum short, acute. Posterior edge of pronotum sinuate. Prosternum slightly longer than prosternal process, process slender, edge curved in lateral view (Figs. 3F, G); procoxal cavities open, narrowly separated (Fig. 3D); promesothoracic clicking mechanism non-functional. Elytra cover abdomen, tapering to apex, widest at humeri, with inconspicuous costae (Fig. 3A). Elytra free apically, independently rounded. Scutellum well developed; widest anteriorly (Fig. 3A) Hind wing well developed.

Legs slightly compressed, with long trochanters (Fig. 3D), tibiae with simple outer edge, bearing setae, and two apical, long spines; tarsomeres slender, five segmented, without ventral pads, claws paired, long, slender, and simple.

Abdomen with six free abdominal sternites, ventrite 1 entire, with inter-coxal process; penultimate tergite simple, ultimate tergite large (Fig. 3P). Male genitalia trilobate; symmetrical. Phallus slender, short (Figs. 3K, M).

Females of all species unknown, putatively incapable of flight.

#### Cebrio igelmimen Rattu et François, 2021

(Fig. 4A)

**Fig. 4.**
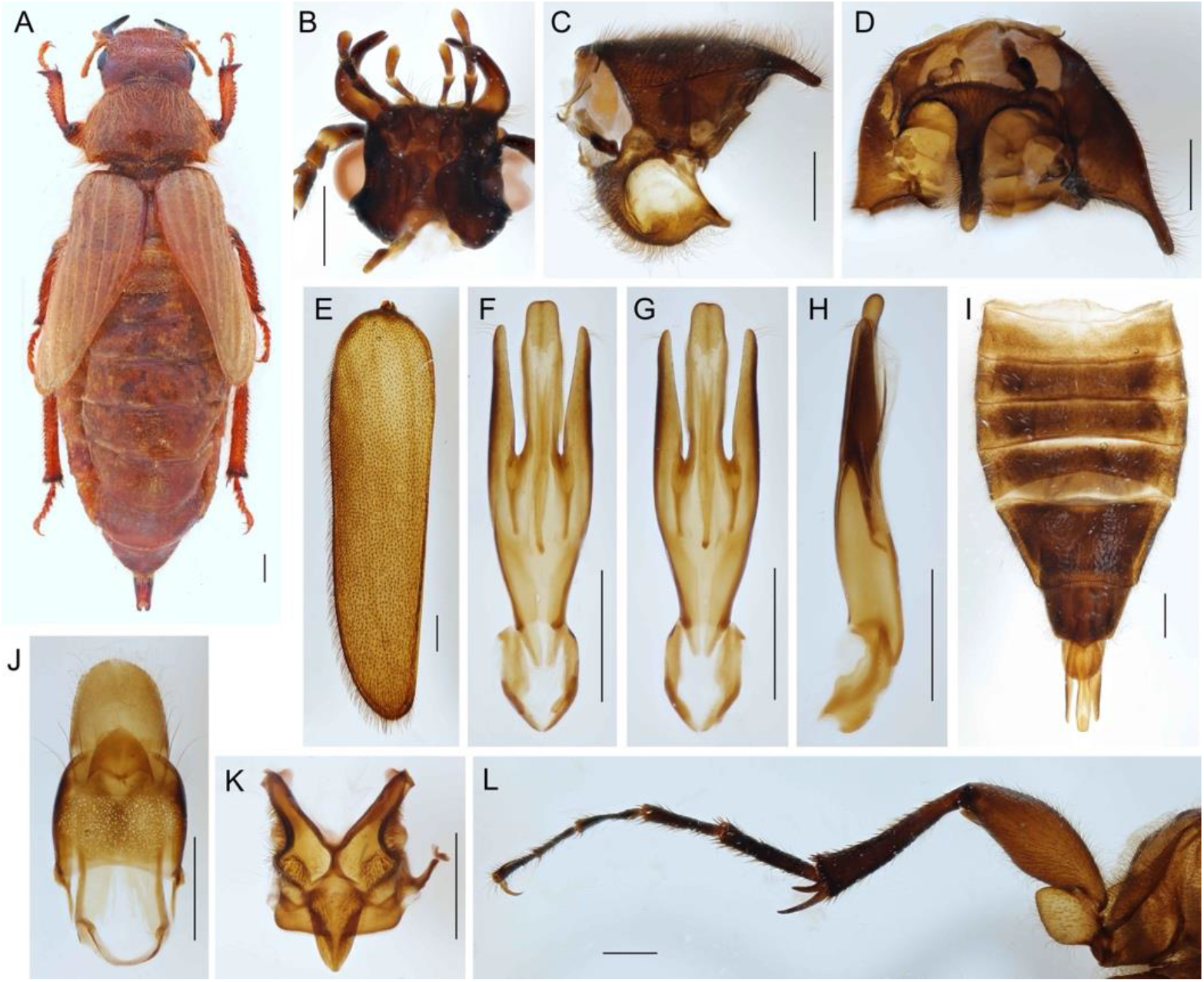
A – *Cebrio (Tibesia) igelmimen* Rattu et François, 2021 from Morocco, female, general appearance. *Cebriorhipis* sp. from Bali. B – head, ventral view; C, D – pronotum, lateral and ventral view; F – elytron; G–I male genitalia, ventral, dorsal and lateral view; J – abdomen, ventral view; K – terminal abdominal segments; L – mesoscutellum; M – metathoracic leg. Scales 1.0 mm. Fig. 4A was published by Rattu & François (2021) and is here reprinted with permission of the authors who retain the copyright of this photo.

#### Remark

The Cebrio females are rare in collections. Recently, two females were described in detail by Rattu (2016) and Rattu & François (2021). Fig. 4A shows the female *Cebrio igelmimen* (photo provided by R. Rattu).

The *Cebrio* females differ from conspecific males in several traits: substantially larger body, small eyes, short antennae, and legs, especially tarsi are substantially shorter than in males (compare Figs. 4A and 4L), often shortened elytra, physogastrous abdomen, weak sclerotization of the cuticle.

#### Cebriorhipis sp

(Fig. 4B–M)

#### Description

Male. Body 12–14 mm long, robust, slightly flattened, brown colored, vestiture dense, long short

(Figs. 4B–E).

Head prognathous to slightly declined, transverse, including eyes narrower than prothoracic width, gradually narrowed posteriorly (Fig. 4B). Eyes slightly protuberant, rounded, ocelli absent. Antennal insertions widely separated, gular sutures narrowly separated, and cervical sclerites sclerotized (Fig. 4B). Antennae slender, shortly flabellate, reaching almost mid of elytra, antennomere 1 robust, long, antennomeres 2 and 3 short, antennomeres 4–11 with short lamella, terminal antennomere similar length as preceding one. Mandibles robust, abruptly curved (Fig. 4E). Mandibular apex and incisor edge simple, maxilla with short setose mala; maxillary palpi slender, 4-segmented, apical palpomere parallel-sided; labium tiny, labial palpi 3-segmented (Fig. 4B).

Prothorax transverse, pronotum without carinae, maximum width 1.5 times length, widest at posterior angles, sides convex (Figs. 4C, D). Prothorax basally narrower than elytral bases; lateral pronotal edge conspicuous in whole length (Fig. 4C). Posterior angles of pronotum ling, acute. Prosternal process four times longer than prosternum (Fig. 5D); procoxal cavities open, narrowly separated; promesothoracic clicking mechanism non-functional (Fig. 4D). Elytra cover abdomen, tapering to apex, widest at humeri, with inconspicuous costae (Fig. 4E). Elytral suture almost complete. Scutellum well developed; pointed posteriorly (Fig. 4K) Hind wings well developed.

**Fig. 5.**
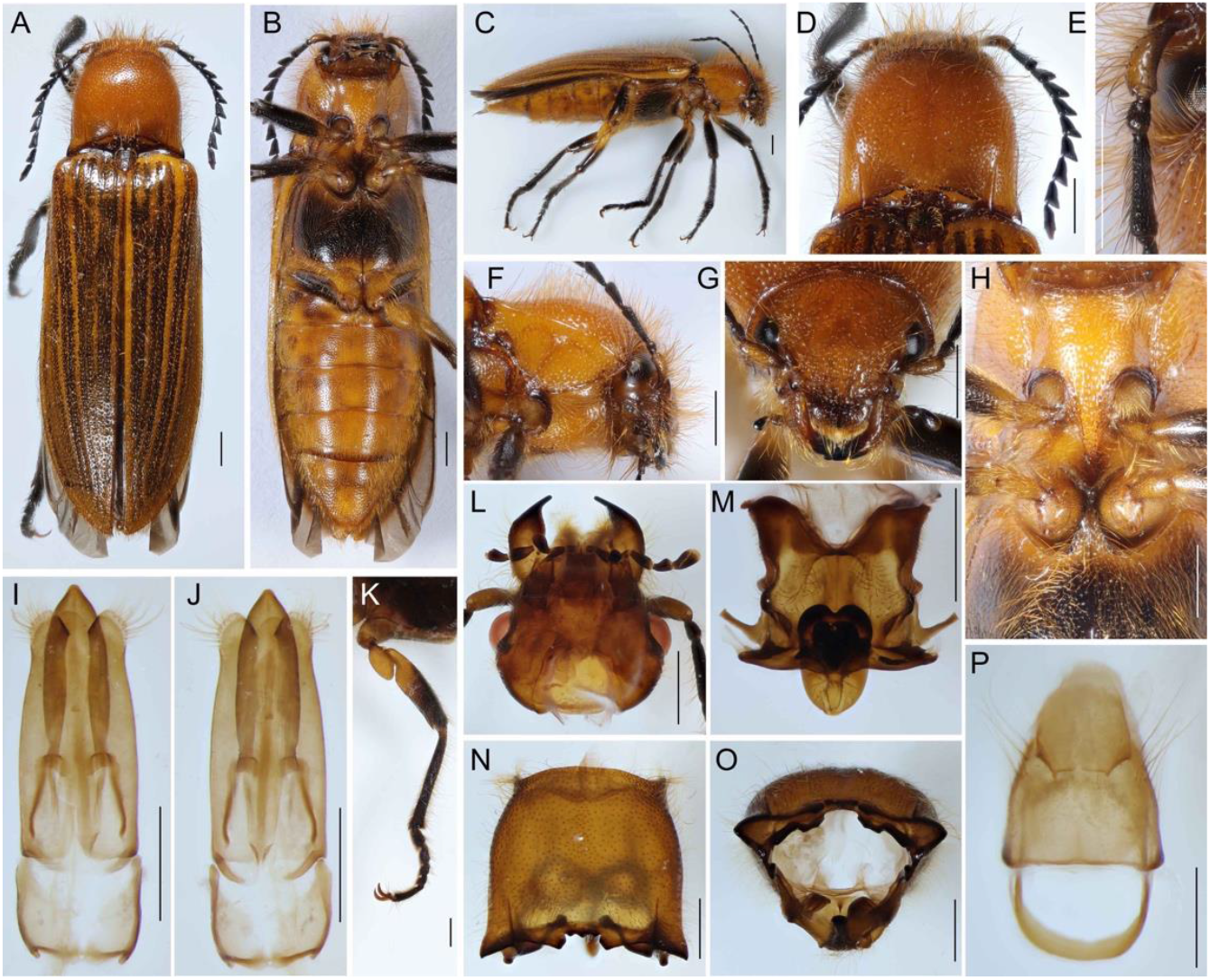
*Hemiopus* sp. from Laos. A–C general appearance, dorsal, ventral, and lateral aspects; D – pronotum; E – basal antennomeres; F – pronotum and head, ventrolateral view; G – head, dorsally; H prosternal process and mesosternal pit; I, J – male genitalia; K – metathoracic leg; L – head ventrally; M – mesonotum; N, O° pronotum, dorsal, and posterior view; P – terminal abdominal segments. Scales 1.0 mm.

Legs slightly compressed, with tarsi longer than femora and tibiae; slender pro- and mesotrochanters and widened metathoracic trochanters (Fig. 4L), tibiae with simple outer edge bearing setae, and two apical, long spines; tarsomeres slender, five segmented, without ventral pads, claws paired, long, slender, and simple (Fig. 4L).

Abdomen with seven free abdominal sternites, ventrite 1 entire, without inter-coxal process; penultimate tergite simple (Fig. 4J). Male genitalia trilobate; symmetrical. Phallus robust, basally fused with parameres (Figs. 4F–H). Females of *Cebriorhipis* unknown.

### Taxonomy

#### Paulusiellinae Kusy, Motyka & Bocak, new subfamily

Type genus

*Paulusiella* Löbl, 2007 (monotypic).

=*Paulusiella* Mandl, 1974 (invalid name).

#### Diagnosis

The erection of the new subfamily is based on molecular relationships that recovered *Paulusiella* as a sister to *Hemiops* Laporte, 1838 (Elateridae: Hemiopinae; Fig. 1). The subfamily is monogeneric, and the description of *Paulusiella serraticornis* is given above. The morphology is illustrated in Fig. 2 and summarized in Tab. 3.

**Tab. 3.** The overview of elaterid morphological characters in the lineages with modified or unknown females and clicking *Hemiopus* sp.

| Character<br>Taxon | Male |  |  |  |  |  |  | Female |
| --- | --- | --- | --- | --- | --- | --- | --- | --- |
|  | anten-<br>nae | apical<br>antenna-<br>meres | prosternum/<br>prosternal<br>process | lateral<br>pronotal<br>edge | elytral rows or<br>costae / suture;<br>shape | abdominal<br>segments | mesocox.<br>process<br>(ventr. 1) | Differences when compared with<br>conspecific male |
| *Hemiops*<br>(Hemiopinae) | short | 10<11 | long / long,<br>clicking | present | present / full; widest<br>in middle | connate | present | fully sclerotized, flying, similar<br>male |
| *Cebrio*<br>(Elaterinae) | short | 10<11 | short / long,<br>non-clicking | present | absent / divergent,<br>widest basally | free | absent | some females with short elytra, and<br>wingless; physogastric abdomen,<br>short antennae, and legs (Fig. 4A). |
| *Paulusiella*<br>(Paulusiellinae) | short | 10<11 | short / short,<br>non-clicking | vestige<br>posteriorly | absent / divergent,<br>widest basally | free | absent | unknown |
| *Analestesa*<br>(Cardiophorinae) | long | 10>11 | long / short,<br>non-clicking | absent | absent / absent;<br>widest basally | free | present | unknown |
| *Plastocerus*<br>(Dendrometrinae) | long | 10<11 | long / short,<br>non-clicking | present | present / full;<br>parallel-sided | free | short | obtuse posterior pronotal angles,<br>shorter antennae than male |
| *Drilus*<br>(Agrypninae) | short | 10<11 | short / short,<br>non-clicking | present | absent / full suture;<br>widest posteriorly | free | absent | head and appendages adult-like;<br>rest of body larviform |
| *Omalisus*<br>(Omalisinae) | long | 10<11 | short / short,<br>non-clicking | present | present; full suture,<br>parallel-sided | free | absent | pupa-like: short legs and antennae,<br>vestigial elytra, similar length of<br>meso- and metathorax |

#### Justification of the erection of the subfamily Paulusiellinae

The morphology does not provide sufficient guidance for the placement of the genus in a natural classification. The absence of apparent diagnostic characters is documented by the initial invalid description of *Paulusiella* in Dascilloidea (Mandl 1974, 1979), the inclusion of an originally karumid species in the genus (*Escalerina serraticornis* Paulus, 1972; Dascilloidea: Dascillidae), and the recent description of a *Paulusiella* species in *Selasia* (*Selasia sweihana* Geisthardt, 2009; Elateridae: Agrypninae: Drilini; Geisthardt 2009; Ivie & Branham 2011). The type-genus *Paulusiella* was validly erected in Cebrioninae (now Elaterinae: Cebrionini) by Löbl (2007) and later transferred to Elateridae *incertae sedis* (Ivie & Barclay 2011).

Here, *Paulusiella* was recovered with robust support as a sister to *Hemiops* Laporte, 1838 by all analyses (Fig. 1A, B, 5A–P, Supp. Figs. S2–S11). Hemiopinae is a small elaterid subfamily with only four genera – *Hemiops* Laporte, 1838 (Figs. 5A–P; East and Southeast Asia), *Legna* Walker, 1858 (Sri Lanka), *Parhemiops* Candèze, 1878 (Nepal) and morphologically somewhat distant *Exoeolus* Broun, 1893 (New Zealand) (Douglas 2011). Comparing *Paulusiella* and *Hemiops*, we can see similar abruptly elevated scutellum that is anteriorly straight in *Paulusiella* but bilobate in *Hemiops* (Figs. 2H, 5M) and similar morphology of trochanters (Figs. 2D, 5B, K; Paulus 1972, Fig. 3). In both taxa, the scutellum posteriorly surpasses the elytral surface, but projected posterior part of the scutellum is commonly encountered in elateroids. The antennae are different, but the relative length of the three basal antennomeres is similar (Figs. 2D, 5D). There are several structures that are present in *Hemiops* and absent in *Paulusiella*: the sharp lateral prothoracic edge (Figs. 2K, 5F), apparent elytral longitudinal costae (Figs. 2I, 5A), the complex posterior shape of the prothorax (Figs. 2J, M, 5N, O). The taxa also differ in the relative length of the prosternal process, prosternum, shape of elytra, abdominal terminal segments, and the shape of phallus (Figs. 2A–Q; 5A–P).

We prefer to assign the subfamily rank to the *Paulusiella*-based taxon as we cannot propose any reliable diagnostic character that would morphologically define the clade of all hemiopine genera and *Paulusiella*.

**Subfamily Cardiophorinae Candèze, 1859**

Cardiophorites Candèze, 1859: 4.

Cardiophori: LeConte 1861: 166.

Cardiophorinae: Burakowski et al. 1985: 227.

Type genus

*Cardiophorus* Eschscholtz, 1829.

=Aphrici LeConte, 1861: 173.

Type genus: *Aphricus* LeConte, 1853.

=Aptopina Jakobson, 1913: 760.

Type genus: *Aptopus* Eschscholtz, 1829.

=Esthesopinae Fleutiaux, 1919: 76.

Type genus: *Esthesopus* Eschscholtz, 1829.

=Nyctorini Semenov-Tian-Shanskij et Pjatakova, 1936: 102.

Type genus: *Nyctor* Semenov-Tian-Shanskij et Pjatakova, 1936 [=*Cardiophorus* Eschscholtz, 1829 *sensu* Cate (2007) but not Douglas (2017)].

=Cebriognathinae Paulus, 1981: 264 (recently Elaterinae: Cebrionini or as a synonym of Cebrioninae), **a new synonym of Cardiophorinae**.

Type genus: *Cebriognathus* Chobaut, 1899 (=*Analestesa* Leach, 1824).

#### Remark

The molecular analysis robustly recovered *Analestesa arabica* in close relationships with *Globothorax* Fleutiaux, 1891 (=*Teslasena* Fleutiaux, 1892), *Cardiophorus* Eschscholtz, 1829, and *Dicronychus* Brullé, 1832 (Figs. 1A, B, Supp. Figs. S2–S11). The position is also supported by the structure of male genitalia (Figs. 3K–M). Although Paulus (1981) erected Cebriognathinae within that time accepted Cebrionidae, he mentioned the here confirmed similarity of the male genitalia of *Cebriognathus* and Cardiophorinae and discussed the possibility that *Cebriognathus* is a modified click beetle. Bouchard et al. (2011) listed Cebriognathinae as a synonym of Cebrioninae, and the GBIF database (https://www.gbif.org/species/4428817 accessed on Nov. 7, 2022) lists *Analestesa* as a junior synonym of *Cebrio*. Still, Analestesa is a valid name and must be considered one of the Cardiophorinae genera.

*Analestesa* is much less sclerotized than most cardiophorine click beetles and has no clicking mechanism (Figs. 3A, D). It differs from most Cardiophorinae in a quadrate mesoscutellar shield and the small and short prosternal process that does not reach the mesothoracic pit. The only cardiophorine species with weakly sclerotized cuticle has been *Nyctor expallidus* Semenov-Tian-Shanskij et Pjatakova, 1936 (=*Cardiophorus expallidus, sensu* Cate 2007). The males have possibly slender antennae, and the females shortened, robust antennae. Although Douglas (2017) described both illustrated specimens as males, the intraspecific variability in the morphology of antennae is improbable (Douglas 2017; Figs. 60, 61).

There are several characters defining Cardiophorinae + Negastriinae and the monophyly of Cardiophorinae (Douglas 2017) that could be sought in *Analestesa*: the shape of the parameres (Figs. 3K–M) and the straight lateral edge of the prosternum. Another diagnostic character was defined in female genitalia, but females are unknown for *Analestesa*. The modified external morphology does not provide evidence for the placement of *Analestesa* in Cardiophorinae, and only the similar male genitalia support the recovered DNA-based topology. The recovered relationships (*Globothorax*(*Analestesa*(*Cardiophorus, Dicronychus*) suggest that the earlier proposed morphology-based topology (*Dicronychus*(*Cardiophorus, Globothorax*) might need further investigation (Douglas 2017).

#### Subfamily Elaterinae Leach, 1815

Elaterides Leach, 1815: 85.

Type genus: *Elater* Linnaeus, 1758.

#### Cebrionini Latreille, 1802

Cebrionates Latreille, 1802: 97.

Type genus: *Cebrio* Olivier, 1790.

Remark

Females of *Cebrio* have variably developed elytra and large bodies. The known females are flightless (Fig. 5A; Rattu 2016, Rattu & François, 2021, Martinez-Luque et al. 2022). The females differ from conspecific males in smaller eyes and a different shape of the cranium, shorter appendages: very short antennae, with low differentiation between the antennomeres, short legs, with longer femora and tibiae than tarsi, shortened elytra, and vestigial wings. *Cebrio gigas* (F., 1787) has a physogastric female but the elytra that form complete elytral suture, although they do not completely cover the abdomen

(https://inpn.mnhn.fr/espece/cd_nom/240515 – accessed on 3 Nov 2022).

#### Revised composition of Cebrionini

Cebrionina Latreille, 1802: *Cebrio* Olivier, 1790; *Cebriorhipis* Chevrolat, 1875, *Musopsis* Chevrolat, 1875, *Scaptolenus* LeConte, 1853; *Selonodon* Latreille, 1834; *Stenocebrio* Solervicens, 1988. Aplastina Stibick, 1979: *Aplastus* LeConte, 1859; *Euthysanius* LeConte, 1853; *Octinodes* Candèze, 1863 (=*Plastocerus* LeConte, 1853); *Cylindroderus* Latreille, 1834 (=*Cylindroderoides* Schwarz, 1907); *Dodecacius* Schwarz, 1902 (Arnett 1949, Johnson 2002, Sánchez-Ruiz & Löbl 2007, Johnson & Chaboo 2015).

Stibick (1979) listed Pleonomini Semenov et Pjatakova, 1936 as a coordinate taxon of Aplastini in Aplastinae. The group was listed as Pleonominae by Cate (2007) and as the tribe Pleonomini in Dendrometrinae by Bouchard et al. (2011). *Pleonomus* Ménétriés, 1849 has both sexes with fully developed elytra but apparent sexual dimorphism (Reitter 1900) resembling the modifications observed in other soft-bodied elaterids.

## Discussion

### Origins of soft-bodied forms

Some fifteen years ago, the Elateridae was a morphologically homogenous beetle family with few slightly modified, weakly sclerotized taxa concentrated in Cebrionini (Elaterinae; Sánchez-Ruiz & Löbl 2007, Lawrence et al. 2011, Bouchard et al. 2011). Until now, cebrionids have served as a collective taxon for at least partly soft-bodied, non-clicking elaterids with flightless or unknown females (Arnett 1949, Johnson 2002, Rattu 2016, Rattu & François 2021, etc.). Since 2007, several DNA-based studies have targeted soft-bodied elateroids. They have advocated that Drilidae, Omalisidae, and Plastoceridae (Cantharoidea are modified click beetles (Bocakova et al. 2007, Kundrata & Bocak 2011, Bocak et al. 2018, Kusy et al. 2018). Additionally, the relationships between widely defined Elateridae and lampyroid families were suggested by Kusy et al. (2021). Other studies recovered a similar clade but rejected the monophyly of Elateridae (Zhang et al. 2018, McKenna et al. 2019, Douglas et al. 2021).

The earlier phylogenetic hypotheses had a clear evolutionary connotation: the shift leading to flightless, soft-bodiedness, the retention of some larva- or pupa-like characters in adults, and even larviform females of some taxa, was understood as a rare phenomenon. Therefore, almost all elateroid lineages with weak sclerotization, including some taxa with neotenic females, shared a hypothesized common ancestor (the Cantharoidea and cantharoid clade concepts; Crowson 1955, 1972, Branham & Wenzel 2003, Lawrence et al. 2011). The rejection of the cantharoid clade (Bocakova et al. 2007, Sagegami-Oba et al. 2007) and the robust placement of some ‘cantharoid’ groups in Elateridae (Kusy et al. 2019) suggested a different view: the process of metamorphosis is less stable than previously thought, and numerous elaterid groups independently lost the clicking mechanism, are soft-bodied (i.e., cantharoid-like), their known females are physogastric, have short appendages, and short, vestigial or absent elytra (Fig. 4A, Tab. 3).

Here, we studied the position of two taxa, *Paulusiella* and *Analestesa* (=*Cebriognathus* or *Cebrio*), earlier placed in Elateridae *incertae sedis* or Elaterinae: Cebrionini) (Sánchez-Ruiz & Löbl 2007, Ivie & Barclay 2011). Both genera are known only in males, and their females are putatively flightless. These are small-bodied, non-clicking, weakly sclerotized beetles, with shortened and posteriorly slender elytra, without a full-length elytral suture and coadaptation between the lateral elytral margins of the abdomen (Figs. 2, 3). Due to these traits, they are superficially similar to some Cebrionini, but the DNA analysis placed them in very distant positions (Figs. 1A, B, Supp. Figs. S2– S11). *Paulusiella serraticornis* was recovered as a sister to Hemiopinae and *Analestesa arabica* as one of the serial splits in Cardiophorinae. These positions are robustly supported by molecular data (Tab. 2). Conversely, their relationships are hardly supported by morphology (see Results). Still, the comparative morphology neither clearly supports their relationships to *Cebrio* (Figs. 2–4). Cebrionini, i.e., *Cebrio, Cebriorhipis*, and *Scaptolenus*, contain medium-sized beetles with a characteristically robust body, short prosternum, fully developed sharp lateral pronotal margin, and seven segments of the abdomen (Fig. 5; Tab. 3). Small-bodied and weakly sclerotized Aplastini, i.e., *Aplastu*s, *Euthysanius*, and *Octinodes*, are more similar due to small and slender body. Still, all have elaterid-like pronotum with acutely projected posterior angles. *Selonodon* is an elaterid-like cebrionid (Galley 1999), and *Stenocebrio* is similar in general appearance to *Paulusiella* or *Analestesa* (Solervicens 1988), but the genus was unavailable for the study.

Although the divergent morphology is sometimes referred to only as flightlessness and soft-bodiedness, the morphological modifications affect almost all body parts, and similar modifications are known in unrelated lineages (e.g., Paulus 1972, Johnston & Gimmel 2020, Tab. 3). Besides shortened to vestigial elytra and wings, we often notice the shortened antennae and legs; the loss or substantial simplification of complex structures, e.g., the pronotum (keels, lateral edge, shortened prosternum, shortened prosternal process, the loss of acutely projected posterior angles, unfunctional click mechanism), a lower ratio between the length of meta and mesothorax (the female of *Omalisus*; Bocak & Brlik 2008), free abdominal segments, the loss of inter-coxal keel in visible abdominal segment 1, loss of costae of rows of punctures in elytra (Figs. 2, 3, 4; Tab. 3).

The fully sclerotized elaterids have thick-walled thoracic segments, strong muscles, a mesosternal pit, long prosternum, and the elytra are held closed by the mesoscutellar catch. Additionally, the prothorax and mesothorax are coadapted, and pivots and flanges enable precisely defined click action (Ewans 1972). This complex mechanism is lost in all soft-bodied groups. The affected groups do not necessarily have all traits modified (e.g., compare the antennal length of *Analestesa* and *Paulusiella*; Figs. 2D, 3I), but the presence of these characters is often recorded in the males of elateroid taxa for which we have proved or hypothesized the modified, flightless females (Crowson 1972, Cicero 1988, Bocakova et al. 2007, Bocek et al. 2018, Kundrata & Bocak 2019, Kusy et al. 2019, Rosa et al. 2020).

The phylogenetic distribution of non-flying and soft-bodied groups is biased to Elateroidea or Elateriformia, respectively (Gould 1977, McMahon & Hayward 2016). Within click beetles, modified forms are known in Elaterinae: Cebrionini, Omalisinae, Dendrometrinae: Plastocerini, and Agrypninae: Drilini (Bocakova et al. 2007, Kundrata & Bocak 2011, Bocak et al. 2018, Bocek et al. 2018, Kusy et al. 2019). The relationships of *Analestesa* and *Paulusiella* are hypothesized as further two independent origins of these modifications: Paulusiellinae is a sister to Hemiopinae, and *Analestesa* is one of the numerous genera of Cardiophorinae (Figs. 1A, B, Supp. Figs. S2–S11).

Similar phenotypes are known in the lampyroid families. Telegeusinae (Omethidae), which are closely related to Artematopodidae, and Jurasaidae, related to Eucnemidae and Cerophytidae, are further examples of the independent origin of soft-bodied forms (Bocakova et al. 2007, Bocak et al. 2014, Zhang et al. 2018, McKenna et al. 2019, Rosa et al. 2020). An additional case of lost sclerotization was hypothesized in Ptilodactylidae, which now contains one morphologically modified species, earlier the type of Podabrocephalidae (Kundrata et al. 2019). Analogical modifications were also reported in one species of *Anorus* (Dascillidae; Johnston & Gimmel 2020) and other dascillids (Karumiinae; Paulus 1972). We suppose, in agreement with earlier studies, that the morphological modifications result from earlier termination of metamorphosis that strongly affects females but also has some effect on males (Gould 1977, Bocak et al. 2008, McMahon & Hayward 2016, etc.).

### Modified morphology and systematics

The modification caused by incomplete metamorphosis (i.e., the retention of some larval characters and the loss of some derived traits) leads to two kinds of taxonomic misplacements. These beetles are often merged into a single clade as the modifications have a similar effect on unrelated lineages. Then, the morphological analyses using the parsimony criterion prefer these shared characters before fewer, if any, contradicting characters shared with close, yet unmodified, relatives. In such a way, earlier authors defined taxa as Cantharoidea and the cantharoid clade (Crowson 1972, Lawrence et al. 2011), the subfamily Cebrioninae or the tribe Cebrionini (Arnett 1949, Bouchard et al. 2011), the clade of neotenic lineages in net-winged beetles (Kazantsev, 2013), or suggested *Podabrocephalus* Pic, 1913 (Byrrhoidea) as a sister of the cantharoid clade (Lawrence et al. 2011).

Additionally, some taxa were initially placed in distantly related but similarly modified groups. For example, Paulus (1972) described *Paulusiella serraticornis* as *Escalerina serraticornis* in Dascillidae, and Geisthardt (2009) described *P. sweihana* in *Selasia* (Agrypninae: Drilini; Ivie & Barclay 2011).

The unrealistically deep rooting of modified forms is a second kind of misplacement. The loss of derived character states leads to the inference of a deeper position of incompletely metamorphosed groups than those of their fully sclerotized relatives. Consequently, the inappropriately high rank is given to modified lineages. This effect led to the earlier discussion on the ancient origin of neotenics (Crowson 1972, Kazantsev 2005), the proposed sister-relationships of *Selonodon* Latreille (Cebrionini) and other elaterids (Lawrence et al. 2011) or descriptions of elaterid subfamilies Pleonominae (=Pleonomini, Dendrometrinae) and Nyctorini (=Cardiophorinae). The absence of female characters in the analyzed matrices (unknown or sometimes completely larviform females) lowers the number of informative characters coded for neotenic taxa. Then, missing data negatively affect the stability of phylogenies.

The above-described pitfalls cannot be solved by any methodological modification of the morphology-based phylogenetic analyses. The earlier analyses handled many taxa and characters and were correctly conducted by experienced comparative morphologists (Branham & Wenzel 2003, Lawrence et al. 2011). In these groups, we urgently need information unaffected by pedogenetic syndrome. Now, we can access information-rich genetic data and reinvestigate the traditionally accepted relationships. The growing evidence suggests common shifts from the clicking, well-sclerotized elaterids to weakly sclerotized forms with highly modified females (Figs. 1A, B, Supp. Figs. Figs. S2–S11). We can look for similar evolutionary pathways in other groups and test the new hypotheses with even more extensive data in the future. The striking conflict between morphology- and DNA-based relationships of extant lineages also calls for cautious analyses of soft-bodied forms preserved in amber deposits, as their relationships cannot be validated with molecular data.

## Acknowledgments

We are very obliged to Vit Kuban (Brno, Czech Republic), the late Willi Büttiker (Basel, Switzerland), Dirk Ahrens, and the late Sylvia Fabrizzi (Bonn, Germany) for providing the critical specimens for the DNA isolation. We thank R. Rattu (Cagliari, Italy) for permission to use his photograph of a *Cebrio* female (Fig. 5A).

## Authors’ contribution and funding

Conceptualization, L.B., D.K., M.M.; molecular analyses, M.M., D.K.; writing, original draft preparation, D.K. M.M., L.B.; comparative morphology, L.B.; reviewing and editing, all co-authors; visualization, L.B., M.M.; funding acquisition, L.B., M.M., D.K. All authors have read and agreed to the published version of the manuscript. This research was funded by The Czech Science Foundation, grant number 22-35327S (L.B., M.M., and D.K.), and the authors.

## Data availability statement

The analyzed sequences are available in the Mendeley depository. Kusy, Dominik; Motyka, Michal; Bocak, Ladislav (2023), “Data for "Ontogenetic modifications produce similar phenotypes in distantly related click beetles (Coleoptera: Elateridae),” DOI:10.17632/73dmw4czm3.1.

