## supplementary materials for "Ontogenetic modifications produce similar phenotypes in distantly related click beetles (Coleoptera: Elateridae)"

### Supplementary Material

**Supplementary Tables**

**Supplementary Table S1.** The earlier sequenced samples and publicly available mitogenomes included in the analyses.

**Supplementary Figures**

**Supplementary Figure S1.** AliStat heat maps of pairwise completeness scores (Ca) in datasets A and D.

**Supplementary Figure S2.** The result of the IQTREE analysis using 15 genes at nucleotide level, partitioned matrix.

**Supplementary Figure S3.** The result of the IQTREE analysis using 15 genes at nucleotide level, unpartitioned matrix.

**Supplementary Figure S4.** The result of the PhyloBayes analysis using 15 genes at nucleotide level.

**Supplementary Figure S5.** The result of the IQTREE analysis using 13 protein coding genes at nucleotide level, partitioned matrix.

**Supplementary Figure S6.** The result of the IQTREE analysis using 13 protein coding genes at nucleotide level, unpartitioned matrix.

**Supplementary Figure S7.** The result of the IQTREE analysis using 13 protein coding genes at nucleotide level masked by Degen, partitioned matrix.

**Supplementary Figure S8.** The result of the IQTREE analysis using 13 protein coding genes at nucleotide level masked by Degen, unpartitioned matrix.

**Supplementary Figure S9.** The result of the IQTREE analysis using 13 protein coding genes at amino acid level, partitioned matrix.

**Supplementary Figure S10.** The result of the IQTREE analysis using 13 protein coding genes at amino acid level, unpartitioned matrix.

**Supplementary Figure S11.** The result of the Phylobayes analysis using 13 protein coding genes at amino acid level.

**Supplementary Table S1.** The earlier sequenced samples and publicly available mitogenomes included in the analyses.

| Taxon | Depository | Acc. # / Voucher | Geogr. origin | Reference / Link to the depository |
|---|---|---|---|---|
| *Sinopyrophorus schimmeli* | GenBank | MH065615 | China: Yunnan | He, J.W., Bi, W., Dong, Z. et al. (2019) The mitochondrial genome of the first luminous click-beetle (Coleoptera: Elateridae) recorded in Asia. Mitochondrial DNA Part B, 4, 565–567. |
| *Agriotes obscurus* | GenBank | KT876879 | unknown | unpublished |
| *Agriotes ustulatus* | GenBank | JX412737 | unknown | unpublished |
| *Adrastus rachifer* | GenBank | KX087232 | unknown | unpublished |
| *Ludioschema vittiger* | GenBank | MN306531 | China: Hainan | Zhao,Y., Wang,Y. and Liu,Y. 2019.The complete mitochondrial genome of click beetle Chiagosnius vittiger (Coleoptera: Elateridae) and phylogenetic analysis. Mitochondrial DNA B Resour 4 (2), 3340-3341 |
| *Elateridae sp.* | GenBank | MH789726 | Borneo | unpublished |
| *Melanotus villosus* | GenBank | KT876904 | unknown | unpublished |
| *Pyrearinus termitilluminans* | GenBank | KJ922150 | Brazil | Amaral, D. T., Mitani, Y., Ohmiya, Y., Viviani, V.R. 2016. Organization and comparative analysis of the mitochondrial genomes of bioluminescent Elateroidea (Coleoptera: Polyphaga) Gene 586, 254-262. |
| *Pyrophorus divergens* | GenBank | EF398270 | Brazil | Arnoldi, F. G. C., Ogoh, K., Ohmiya, Y., Viviani, V. R. 2007. Mitochondrial genome sequence of the Brazilian luminescent click beetle Pyrophorus divergens (Coleoptera: Elateridae): mitochondrial genes utility to investigate the evolutionary history of Coleoptera and its bioluminescence<br>Gene, 405, 1-9. |
| *Ignelater luminosus* | GenBank | MG242621 | Brazil | Amaral, D. T., Mitani, Y., Ohmiya, Y., Viviani, V. R. 2016. Organization and comparative analysis of the mitochondrial genomes of bioluminescent Elateroidea (Coleoptera: Polyphaga) Gene 586, 254-262. |
| *Hapsodrilus ignifer* | GenBank | KJ922149 | Brazil | Amaral, D. T., Mitani, Y., Ohmiya, Y., Viviani, V. R. 2016. Organization and comparative analysis of the mitochondrial genomes of bioluminescent Elateroidea (Coleoptera: Polyphaga) Gene 586, 254-262. |
| *Cryptalaus yamato* | GenBank | MK524933 | Korea | Lee, S.-G., Choi, S., Roh, S. J., Lee, B.-W. and Lim, J. 2019. A first record of complete mitochondrial genome of Cryptalaus Ohira (Insecta: Coleoptera: Elateridae) based on C. yamato (Nakane) Mitochondrial DNA B Resour 4 (1), 1628-1629. |
| *Cryptalaus larvatus* | GenBank | MT118665 | unknown | unpublished |

| | | | | |
|---|---|---|---|---|
| *Agrypnus sp.* | GenBank | MN370897 | Korea, Gyeonggi Prov. | Du,Y., Liu,X., Long,Y., Cheng,S. and Zhong,B. The complete mitochondrial genome of click beetle Agrypnus sp. (Coleoptera: Elateridae) and phylogenetic analysis. Mitochondrial DNA B Resour 4 (2), 3354-3355 (2019) |
| *Drilus flavescens* | GenBank | HQ232815 | Malta | Timmermans, M. J., Dodsworth,S., Culverwell, C. L., Bocak, L., Ahrens, D., Littlewood, D. T., Pons, J. and Vogler, A. P. Why barcode? High-throughput multiplex sequencing of mitochondrial genomes for molecular systematics. Nucleic Acids Res. 38 (21), E197 (2010) |
| *Anostirus castaneus* | GenBank | KX087237 | unknown | unpublished |
| *Limonius minutus* | GenBank | KX087306 | unknown | unpublished |
| *Limonius californicus* | GenBank | KT852377 | USA | Gerritsen, A. T., New, D. D., Robison, B.D., Rashed, A., Hohenlohe, P.,Forney,L., Rashidi, M., Wilson, C. M. and Settles, M. L. 2016. Full Mitochondrial Genome Sequence of the Sugar Beet Wireworm Limonius californicus (Coleoptera: Elateridae), a Common Agricultural Pest. Genome Announc 4 (1), e01628-15. |
| *Athous haemorrhoidalis* | GenBank | KT876881 | unknown | Linard, B., Arribas, P., Andujar, C., Crampton-Platt, A. and Vogler, A. P. Lessons from genome skimming of arthropod-preserving ethanol Mol Ecol Resour (2016). |
| *Cardiophorus signatus* | GenBank | MK692585 | Spain | Andujar, C., Arribas, P., Motyka, M., Bocek, M., Bocak, L., Linard,B. and Vogler, A. P. New mitochondrial genomes of 39 soil dwelling Coleoptera from metagenome sequencing. Mitochondrial DNA B Resour 4 (2), 2447-2450 (2019) |
| *Dicronychus cinereus* | GenBank | KX087283 | unknown | unpublished |
| *Dicronychus sp.* | GenBank | JX412848 | unknown | unpublished |
| *Globothorax femoralis* | GenBank | KJ938491 | Brazil | Amaral, D. T., Mitani, Y., Ohmiya, Y., Viviani, V.R. 2016. Organization and comparative analysis of the mitochondrial genomes of bioluminescent Elateroidea (Coleoptera: Polyphaga) Gene 586, 254-262. |
| *Diplophoenicus* sp. | Mendeley | G19011 | Madagascar | Kusy, D.; Motyka, M.; Bocak, L. 2021. Click Beetle Mitogenomics with the Definition of a New Subfamily Hapatesinae from Australasia (Coleoptera: Elateridae). Insects 12, 17.<br>DOI:10.17632/gv88vzb7vn.1 |
| *Drilus mauritanicus* | Mendeley | G18004 | Spain | Kusy, D.; Motyka, M.; Bocak, L. 2021. Click Beetle Mitogenomics with the Definition of a New Subfamily Hapatesinae from Australasia (Coleoptera: Elateridae). Insects 12, 17.<br>DOI:10.17632/gv88vzb7vn.1 |
| *Tibionema abdominalis* | Mendeley | G20012 | Chile | Kusy, D.; Motyka, M.; Bocak, L. 2021. Click Beetle Mitogenomics with the Definition of a New Subfamily Hapatesinae from Australasia (Coleoptera: Elateridae). Insects 12, 17.<br>DOI:10.17632/gv88vzb7vn.1 |
| *Parablax* sp. | Mendeley | G19006 | Queensland | Kusy, D.; Motyka, M.; Bocak, L. 2021. Click Beetle Mitogenomics with the Definition of a New Subfamily Hapatesinae from Australasia (Coleoptera: Elateridae). Insects 12, 17. |

| | | | | |
|---|---|---|---|---|
| | | | | DOI:10.17632/gv88vzb7vn.1 |
| *Eudicronychus rufus* | Mendeley | G19007 | Zambia | Kusy, D.; Motyka, M.; Bocak, L. 2021. Click Beetle Mitogenomics with the Definition of a New Subfamily Hapatesinae from Australasia (Coleoptera: Elateridae). Insects 12, 17. DOI:10.17632/gv88vzb7vn.1 |
| *Eudicronychus* sp. | Mendeley | G20004 | Zambia | Kusy, D.; Motyka, M.; Bocak, L. 2021. Click Beetle Mitogenomics with the Definition of a New Subfamily Hapatesinae from Australasia (Coleoptera: Elateridae). Insects 12, 17. DOI:10.17632/gv88vzb7vn.1 |
| *Hapatesus tropicus* | Mendeley | G20007 | New Guinea | Kusy, D.; Motyka, M.; Bocak, L. 2021. Click Beetle Mitogenomics with the Definition of a New Subfamily Hapatesinae from Australasia (Coleoptera: Elateridae). Insects 12, 17. DOI:10.17632/gv88vzb7vn.1 |
| *Plastocerus angulosus* | Mendeley | A01544 | Turkey | Kusy, D.; Motyka, M.; Bocak, L. 2021. Click Beetle Mitogenomics with the Definition of a New Subfamily Hapatesinae from Australasia (Coleoptera: Elateridae). Insects 12, 17. DOI:10.17632/gv88vzb7vn.1 |

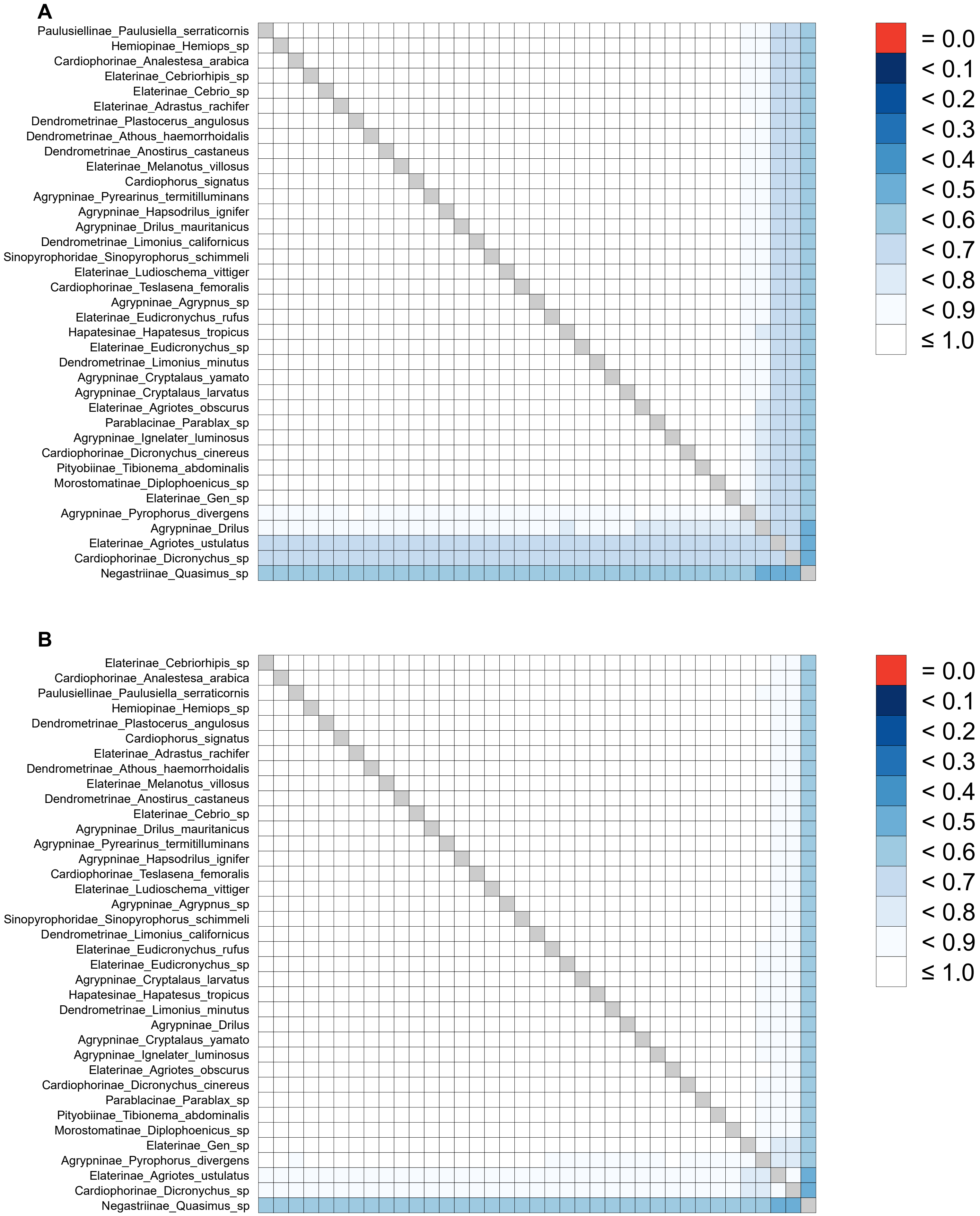


Supplementary Figure S1. AliStat heat maps of pairwise completeness scores (Ca) in datasets A and D:
(A) nucletiode level, 15 mitochondrial genes (2 rRNAs and 13 PGGs, 0.928189),
(D) amino acid level: 13 PCGs (0.952717), analysed using AliScore.

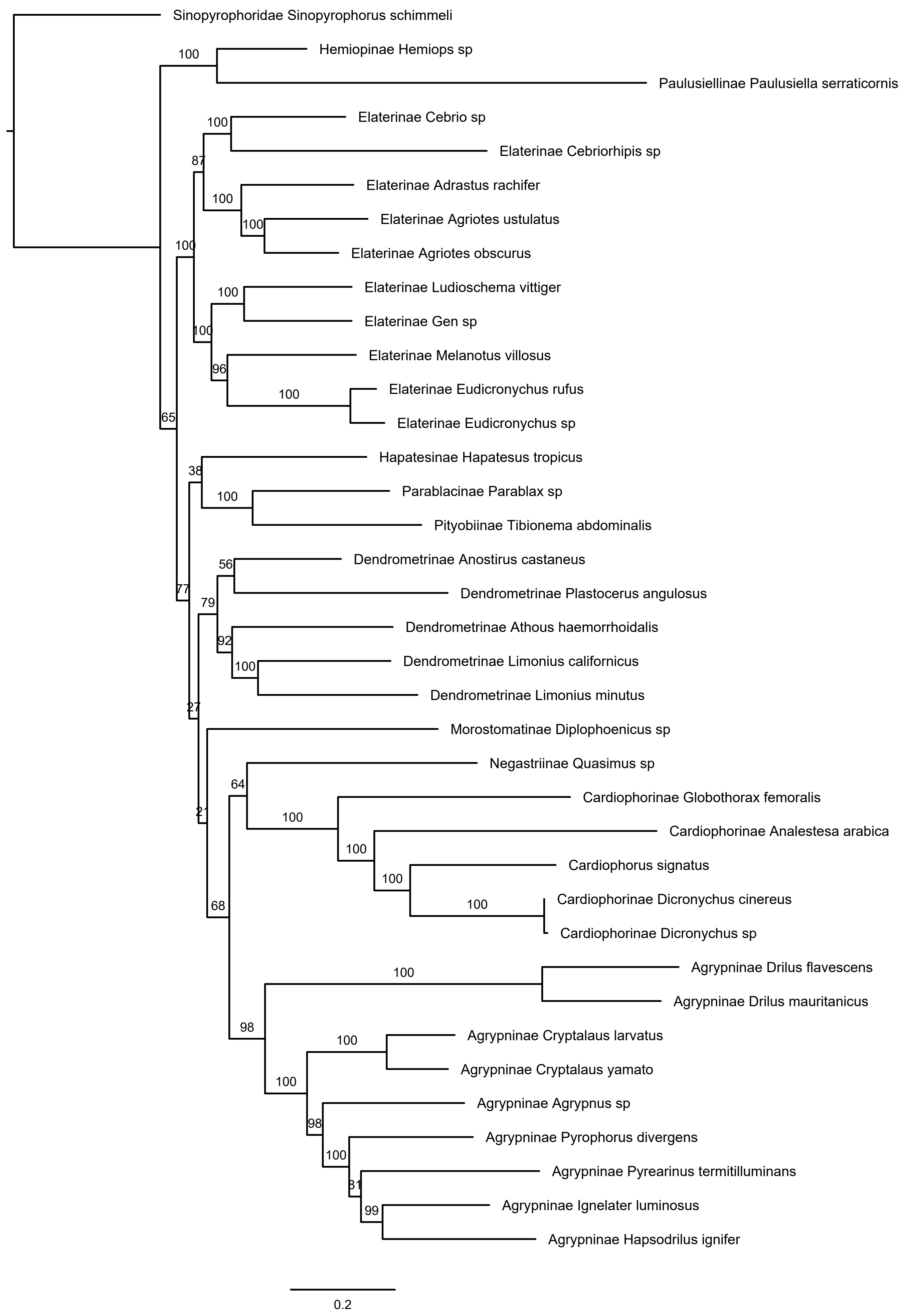


Supplementary Figure S2. The result of the IQTREE analysis using 15 genes at nucleotide level, partitioned matrix.

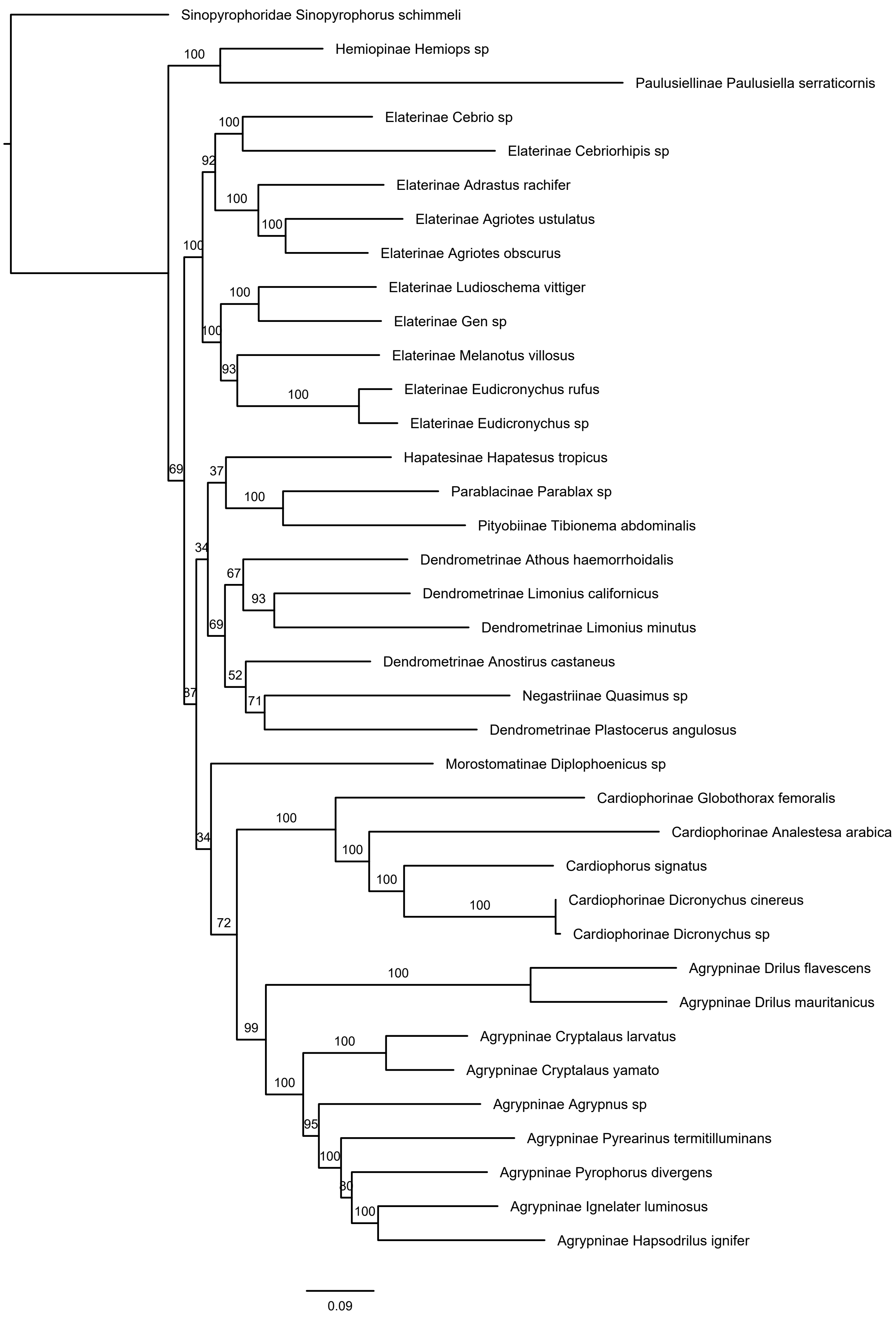


Supplementary Figure S3. The result of the IQTREE analysis using 15 genes at nucleotide level, unpartitioned matrix.

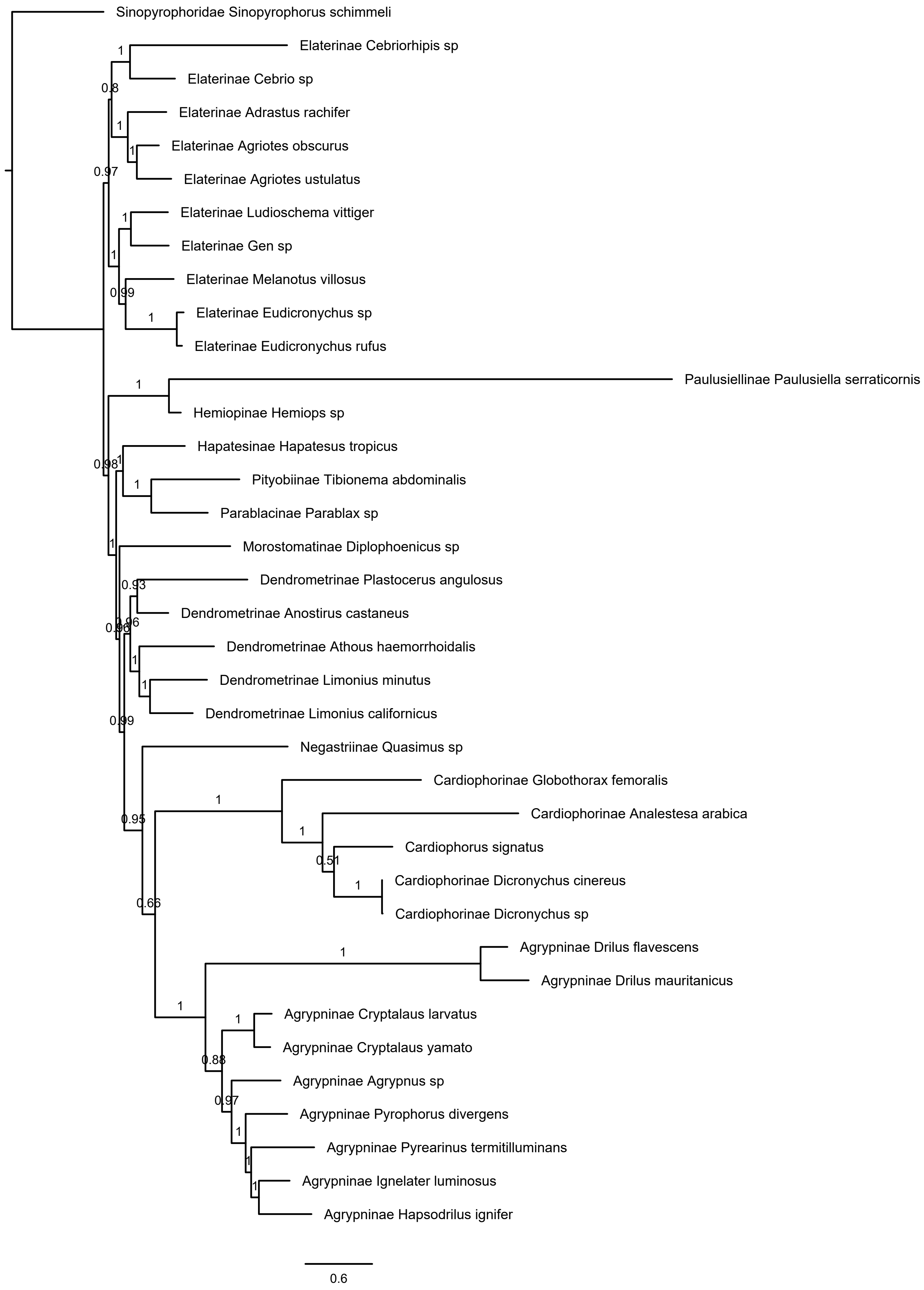


Supplementary Figure S4. The result of the PhyloBayes analysis using 15 genes at nucleotide level.

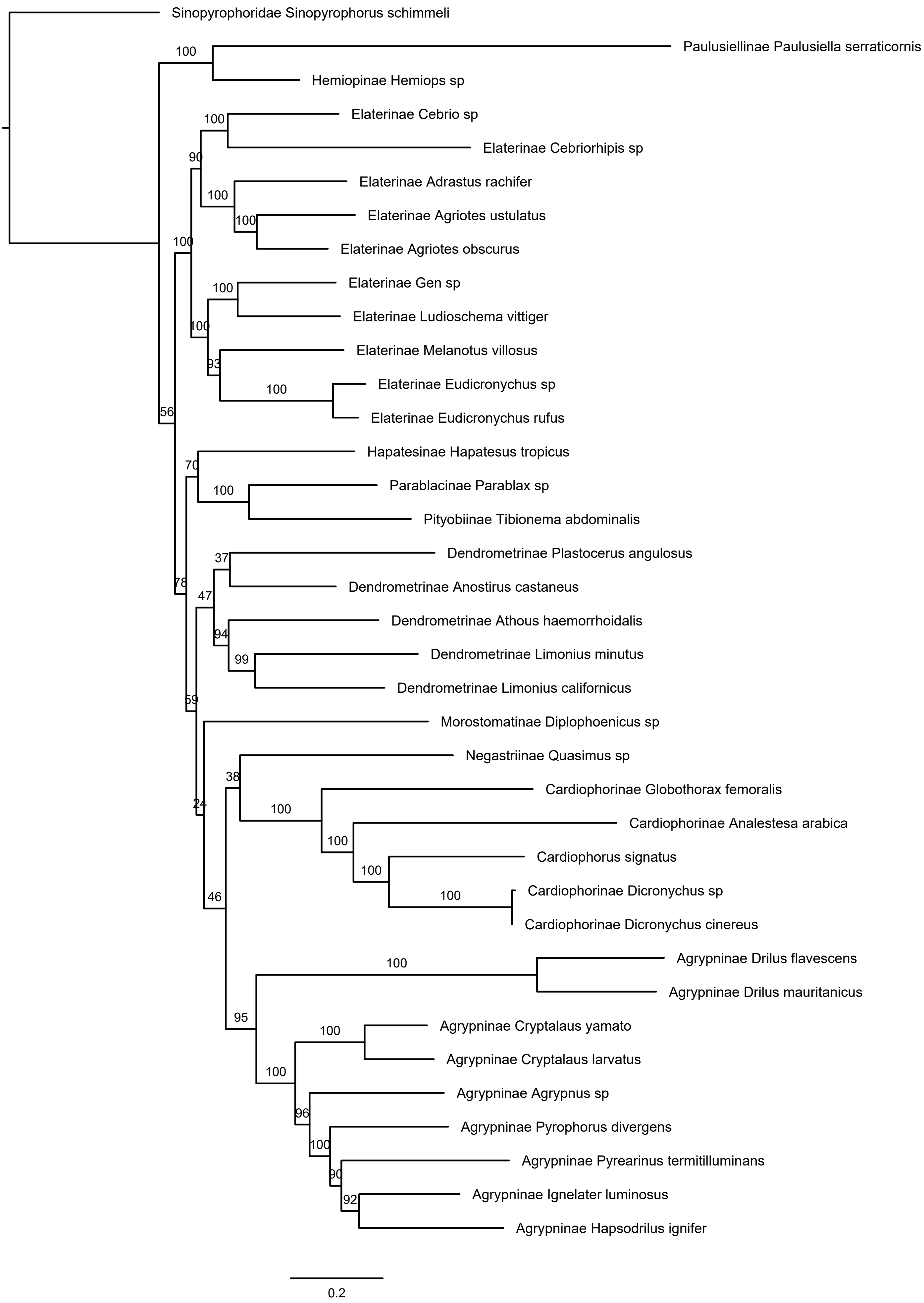


Supplementary Figure S5. The result of the IQTREE analysis using 13 protein coding genes at nucleotide level, partitioned matrix.

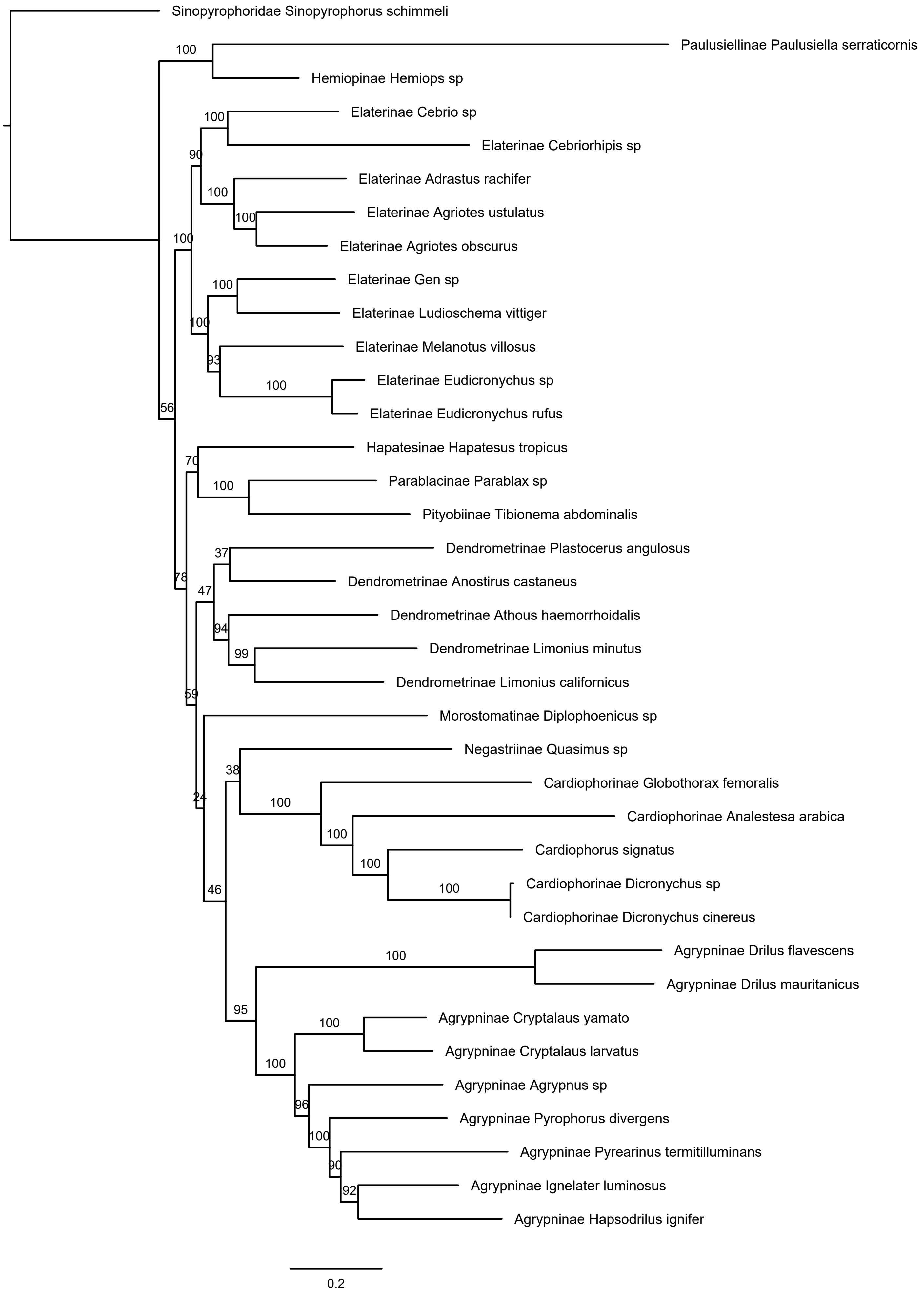


Supplementary Figure S6. The result of the IQTREE analysis using 13 protein coding genes at nucleotide level, unpartitioned matrix.

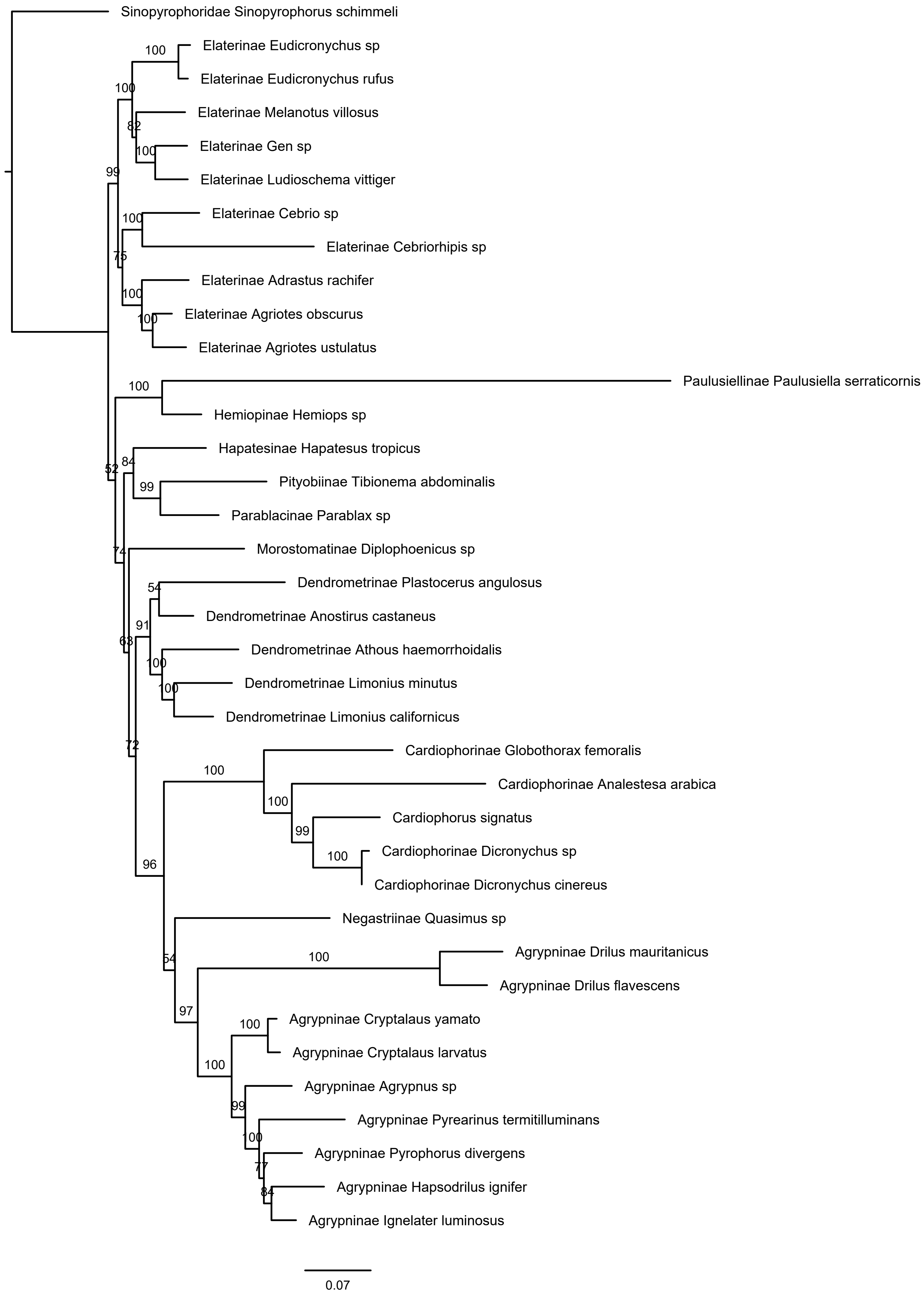


Supplementary Figure S7. The result of the IQTREE analysis using 13 protein coding genes at nucleotide level masked by Degen, partitioned matrix.

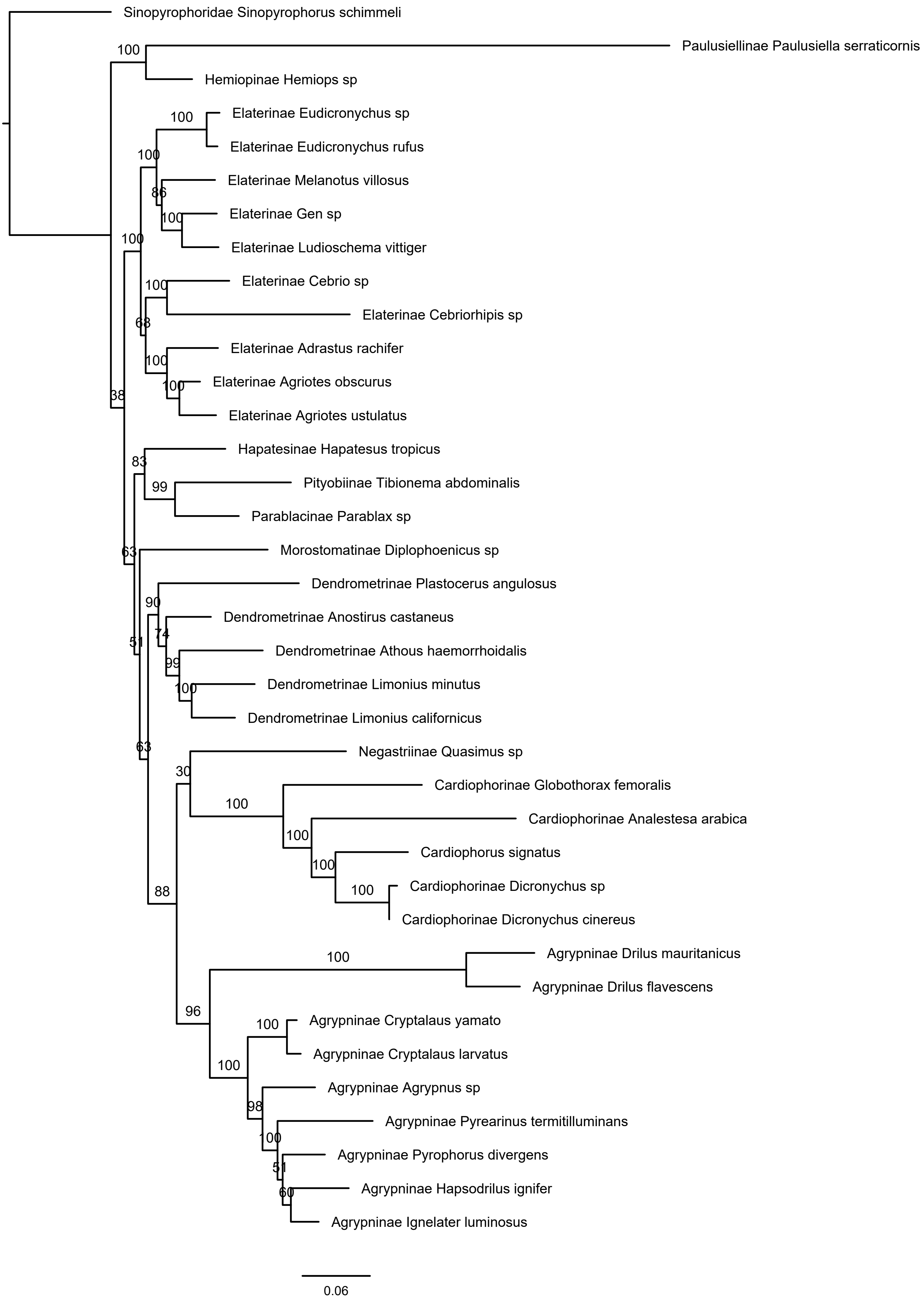


Supplementary Figure S8. The result of the IQTREE analysis using 13 protein coding genes at nucleotide level masked by Degen, unpartitioned matrix.

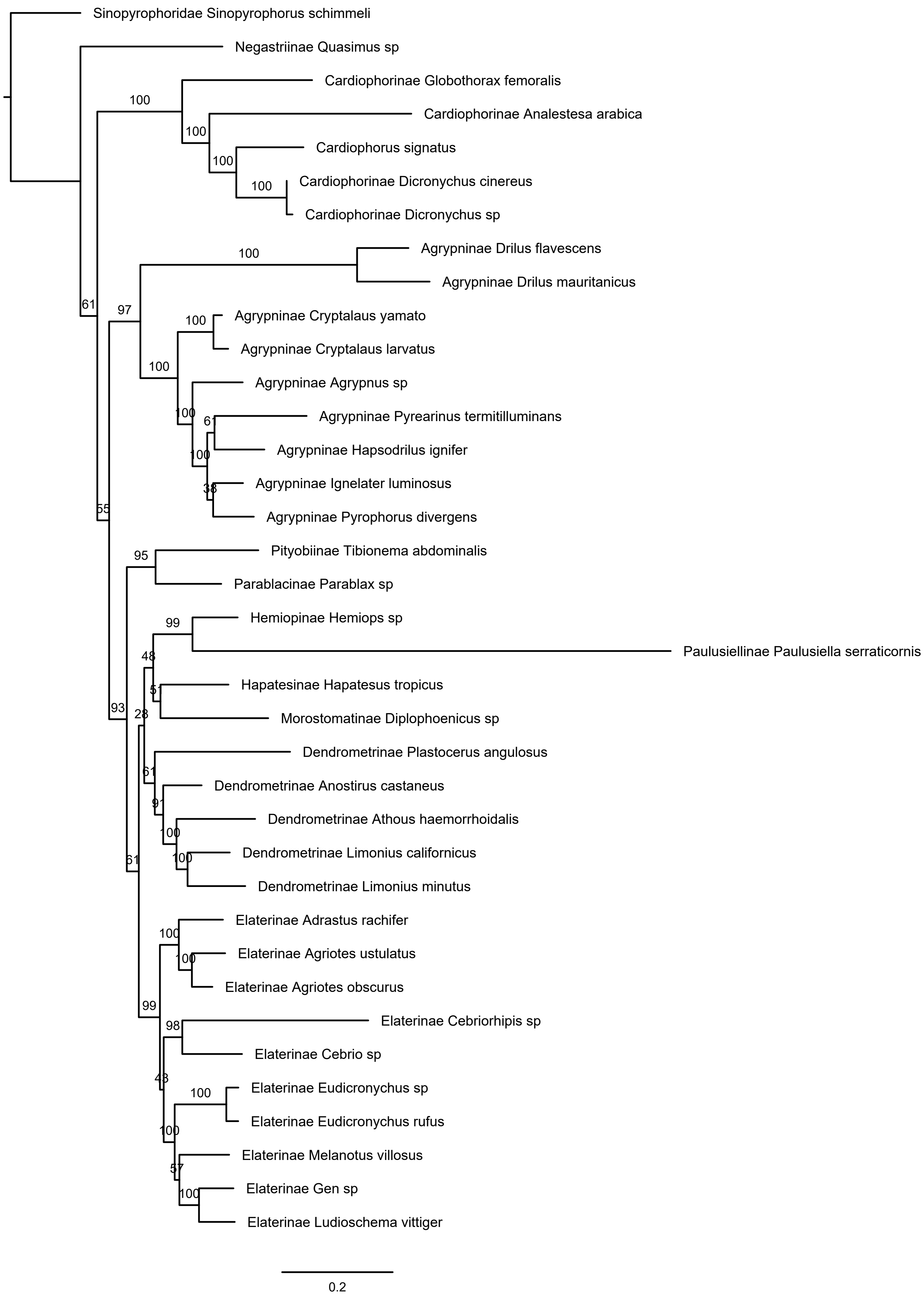


Supplementary Figure S9. The result of the IQTREE analysis using 13 protein coding genes at amino acid level, partitioned matrix.

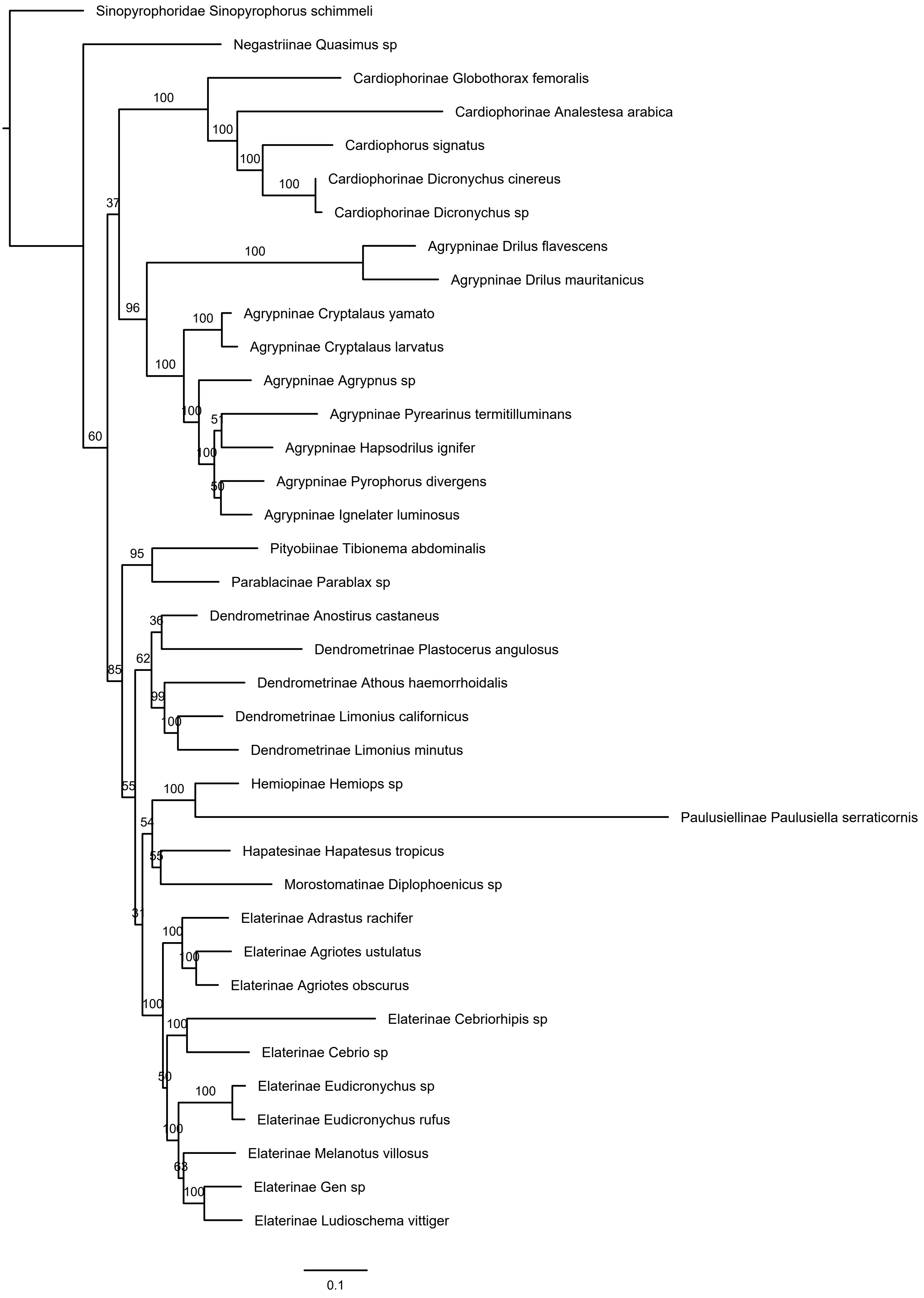


Supplementary Figure S10. The result of the IQTREE analysis using 13 protein coding genes at amino acid level, unpartitioned matrix.

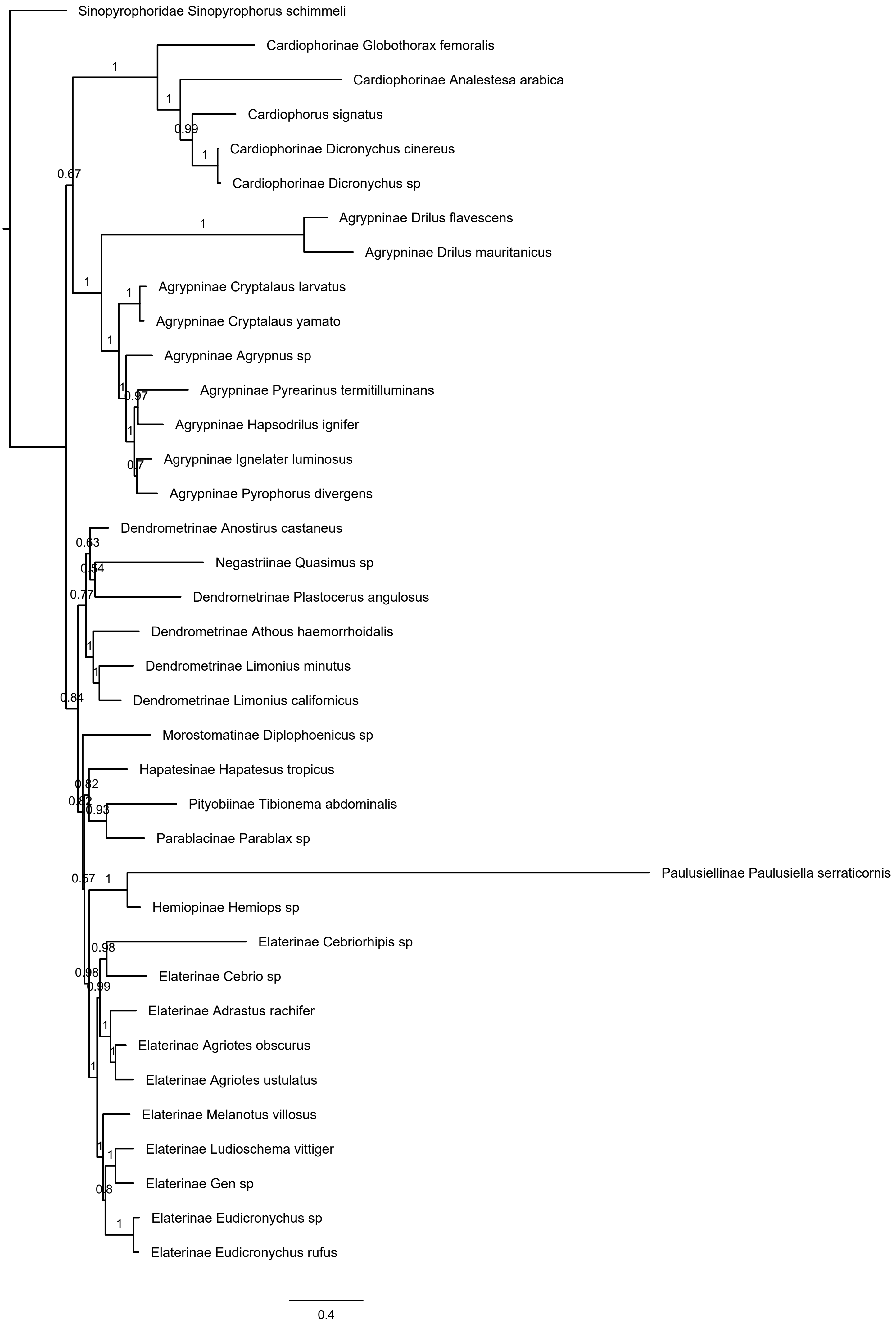


Supplementary Figure S11. The result of the Phylobayes analysis using 13 protein coding genes at amino acid level.
